# Reference-free deconvolution of complex DNA methylation data – a systematic protocol

**DOI:** 10.1101/853150

**Authors:** Michael Scherer, Petr V. Nazarov, Reka Toth, Shashwat Sahay, Tony Kaoma, Valentin Maurer, Christoph Plass, Thomas Lengauer, Jörn Walter, Pavlo Lutsik

**Affiliations:** Department of Genetics/Epigenetics, Saarland University, 66123 Saarbrücken, Germany; Research Group Computational Biology, Max Planck Institute for Informatics, 66123 Saarbrücken, Germany; Graduate School of Computer Science, Saarland Informatics Campus, 66123 Saarbrücken, Germany; Quantitative Biology Unit, Luxembourg Institute of Health, 1445 Strassen, Luxembourg; Division of Cancer Epigenomics, German Cancer Research Center (DKFZ), 69120 Heidelberg, Germany

## Abstract

Epigenomic profiling enables unique insights into human development and diseases. Often the analysis of bulk samples remains the only feasible option for studying complex tissues and organs in large patient cohorts, masking the signatures of important cell populations in convoluted signals. DNA methylomes are highly cell type-specific, and enable recovery of hidden components using advanced computational methods without the need for reference profiles. We propose a three-stage protocol for reference-free deconvolution of DNA methylomes comprising: (i) data preprocessing, confounder adjustment and feature selection, (ii) deconvolution with multiple parameters, and (iii) guided biological inference and validation of deconvolution results. Our protocol simplifies the analysis and integration of DNA methylomes derived from complex samples, including tumors. Applying this protocol to lung cancer methylomes from TCGA revealed components linked to stromal cells, tumor-infiltrating immune cells, and associations with clinical parameters. The protocol takes less than four days to complete and requires basic R skills.

## 1 Introduction

DNA methylation, preferentially occurring at CpG dinucleotides in mammalian genomes, correlates with cell type identity and differentiation stage^1^. Importantly, methylation states of particular CpGs can be used as powerful biomarkers for various conditions, including cancer^2–4^, inflammatory diseases^5^ and aging^6^. To facilitate such findings, large international consortia, including IHEC^7^, DEEP^1^, and BLUEPRINT^8^, are generating genome-wide DNA methylation maps (or DNA methylomes) of primary tissue samples and isolated cell populations. However, DNA methylomes obtained from bulk samples are intrinsically heterogeneous, and can be additionally affected by aging, sex, and technical confounders. Computational methods for the integration of large-scale DNA methylation datasets, and capable of delineating such complex methylomes into biologically distinct components of variability, are of paramount importance^9^.

Deconvolution methods dissect methylomes of cell mixtures into their basic constituents^10^. In case reference DNA methylomes of purified cell types are available, they can be used to infer the proportions of different cell types across the samples. Multiple reference-based methods have been proposed^11–14^ and are reviewed elsewhere^15^ (summarized in **Supplementary Table 1**). Another class of deconvolution methods allows for composition-adjusted identification of differentially methylated regions in epigenome-wide association studies (EWAS) without explicitly computing the cell type proportions^16,17^. When reference methylomes and other prior information is partially or completely absent, semi-reference-free^18^ or fully reference-free^19–23^ deconvolution methods can be applied (**Supplementary Table 1**). Reference-free methods are of particular benefit for poorly characterized, complex systems such as tissues include brain, and solid tumors, where adequate reference profiles are especially hard to obtain. In a separate study, we found that in a large variety of applications, different reference-free deconvolution tools are only marginally different in their accuracy, and a thorough preprocessing, as well as careful feature selection are more important than the choice of the deconvolution tool^25^. Furthermore, interpretation of deconvolution results is challenging in case prior information about the investigated biological system is limited. Here, we present a comprehensive pipeline that facilitates reference-free deconvolution, starting from raw DNA methylation data down to result interpretation. Although we focus on *MeDeCom* as a representative method, the protocol is not limited to a single deconvolution method and can be used in combination with other available tools.

**Table 1:** Quality filtering steps and default parameters in *DecompPipeline*.

| STEP | SITES FILTERED | DEFAULT PARAMETERS |
| --- | --- | --- |
| Bead filtering | Coverage less than *MIN_N_BEADS* beads in any of the samples | *FILTER_BEADS=TRUE*<br>*MIN_N_BEADS=3* |
| Intensity filtering | High or low coverage outliers according to the intensity quantiles specified | *FILTER_INTENSITY=TRUE*<br>*MIN_INT_QUANT=0.001*<br>*MAX_INT_QUANT=0.999* |
| Missing value filtering | Missing measurement in any of the samples | *FILTER_NA=TRUE* |
| SNP filtering | SNPs according to the *dbSNP*<sup>33</sup> database | *FILTER_SNP=TRUE* |
| Somatic site filtering | Located on the sex chromosomes | *FILTER_SOMATIC=TRUE* |
| Cross-reactive filtering | Reported to be cross reactive<sup>34</sup> | *FILTER_CROSS_REACTIVE=TRUE* |

### 1.1 Development of the protocol

Reference-free deconvolution is a computationally demanding task, for which several methods have been proposed, along with the tools implementing them (**Supplementary Table 1**). However, in a pilot benchmark of several published reference-free deconvolution tools, we found that their performance differences were marginal, both on fully synthetic and *in silico* mixed experimental datasets^25^. In fact, the quality and information content of the input DNA methylation matrix had a higher impact upon the accuracy, than the choice of the deconvolution tool. Thus, deconvolution algorithms *a priori* rely on thorough data preprocessing and feature selection, especially if the differences between underlying components are small. Furthermore, biological interpretation of deconvolution results is often challenging, in particular for beginners with limited bioinformatic experience. In order to facilitate analysis and interpretation of complex DNA methylation data, we developed a comprehensive, three-stage protocol, which includes critical preprocessing and interpretation steps in addition to the actual deconvolution. The protocol, schematically outlined in **Fig. 1**, consists of three main stages: (i) meticulous preprocessing of DNA methylation data, including stringent, quality-adapted CpG filtering, elimination of potential confounding factors using Independent Component Analysis (ICA) and feature selection; (ii) reference-free methylome deconvolution for poorly understood biological systems for which no reference profiles are available; (iii) interpretation of deconvolution results with a user-friendly R/Shiny-based interface, enabling generating novel biological insights.

**Fig. 1:**
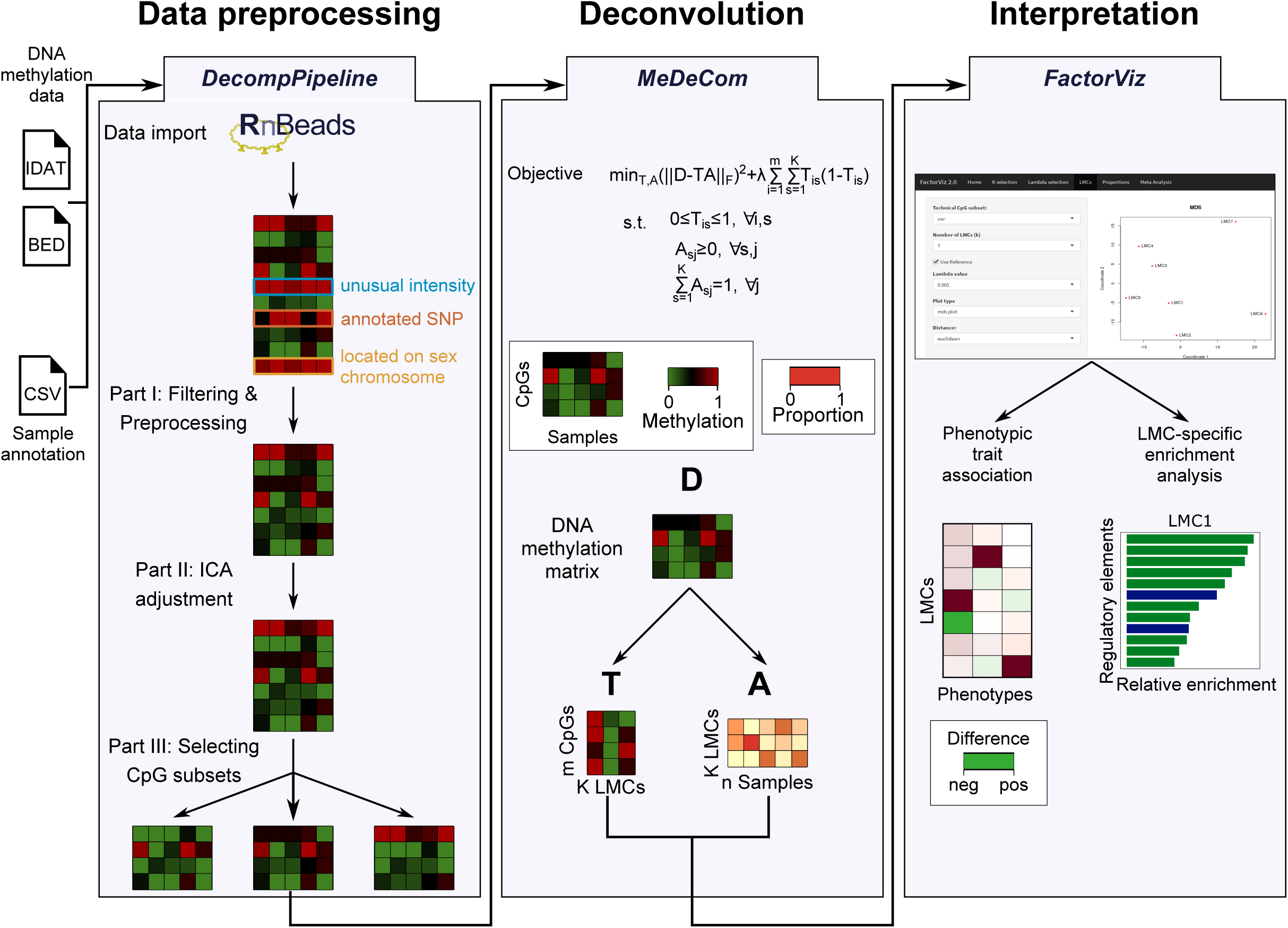
Overview of the proposed deconvolution protocol. DNA methylation data can be used from any technology yielding single CpG methylation calls. Methylation data is first processed using *DecompPipeline*, which includes data import, preprocessing, accounting for confounders and feature selection. *MeDeCom* can be used to perform deconvolution of the input methylation matrix (dimension *m* CpGs x *n* samples) into the latent methylation components (LMCs) and the proportions matrix (dimension *K* LMCs x *n* samples), while the protocol is also applicable to different deconvolution tools. The resulting matrices are then validated and interpreted using the R/Shiny visualization tool *FactorViz*.

Data preprocessing is key to the overall success of deconvolution. The first stage of our protocol thus comprises removal of unreliable or otherwise problematic measurements using the widely used *RnBeads* software package for handling DNA methylation data^26,27^. Age, sex or donor genotype, can have a strong influence on the methylome, and investigators may want to adjust for those in their analyses^28,29^, i.e. to consider these factors as confounders. Therefore, we argue that accounting for confounding factors, using methods such as Independent Component Analysis (ICA)^30^, is crucial to obtaining biologically relevant results. As the final processing step, a CpG subset selection determines sites that are linked to, for instance, cell type identity or any other phenotypic trait of interest.

The prepared, high-quality data matrix can be subjected to one of the deconvolution tools (**Supplementary Table 1**). These methods decompose the input DNA methylation matrix into a matrix of latent methylation components (LMCs, *T*) and a matrix of proportions (*A*). In the second stage of the protocol, we use *MeDeCom*^21^, our own method based on regularized non-negative matrix factorization (NMF) for this purpose, but the protocol seamlessly integrates with other deconvolution tools.

Although reference-free deconvolution is flexible with respect to the investigated system, the interpretation of deconvolution results, especially of the LMC matrix *T*, can be challenging. In addition to cell type profiles, LMCs reflect multiple drivers of biological and technical variability. Furthermore, validating the proportions and LMCs is not trivial, since the space of possible solutions is large and the cellular composition is typically unknown. In order to guide users, we implemented most of the interpretation functionality as a specialized R/Shiny-based graphical user interface (*FactorViz*).

### 1.2 Applications of the methods

Reference-free deconvolution is the method of choice for studying heterogeneity of DNA methylomes in biological systems with limited prior knowledge about their cellular composition, or in case of missing reference profiles. This includes EWAS using material from hardly accessible or insufficiently characterized organs and tissues, such as human brain, as well as solid tumors. Previously, we and others used this approach to understand cellular heterogeneity in placenta^31^, multiple sclerosis^32^, breast cancer^20^, and cholangiocarcinoma^33^. Reference-free deconvolution is particularly useful to dissect tumor heterogeneity, e.g. to study the effect of tumor-infiltrating immune cells on the tumor microenvironment^34^. Furthermore, identified LMCs can be correlated to tumor size, location, metastasis state, and mutational burden. Since, in general, tumors show a high degree of sample-to-sample variation, methylome deconvolution can be used to detect similarities among different types of cancers to define pan-cancer and cancer type-specific markers. If a particular cancer induces changes in the DNA methylation pattern of the tumor stroma, these changes are likely to be missed by reference-based methods, but not so by reference-free methods.

In the original publication, along with the validation on simulated data and *in-silico* cell type mixtures, *MeDeCom* was applied to a brain frontal cortex dataset, successfully identifying neuronal and glia fractions, as well as detecting additional LMCs, which could be linked to features of Alzheimer’s disease^21^. We anticipate successful application of reference-free deconvolution in similar scenarios. Furthermore, although for blood-based studies reference methylomes exist and reference-based methods perform generally well, reference-free deconvolution can be useful in case of severely altered blood composition, e.g. due to an overproduction of rare cell types. Finally, in the case of blood and other similarly well characterized tissues, *MeDeCom* can be applied in a semi-supervised fashion, i.e. comparing the obtained LMCs to available reference profiles. This enables easy recovery of known signatures, and allows for detection of additional unknown LMCs.

Single-cell technologies are steadily improving, and cell-level DNA methylomes will become increasingly available in the future^35,36^. Nevertheless, we expect that deconvolution of large-scale bulk tissue datasets will remain a useful complement to single-cell DNA methylation profiling, which still suffers from high costs, low sample throughput and data sparsity. We envisage that both approaches can be successfully used in combination, e.g. single-cell profiling for several reference samples in controlled study settings followed by deconvolution of bulk methylomes from large patient cohorts, where single-cell profiles can be used for interpretation of the LMCs. Finally, deconvolution of more accessible bulk methylomes can be used to integrate them with easier to obtain single-cell profiles of other data, including single-cell transcriptomes and chromatin accessibility maps.

### 1.3 Outline of the procedure

Our protocol is divided into three main stages: (i) Data preprocessing, (ii) Deconvolution and (iii) Interpretation, which are described in detail below (see also **Fig. 1**).

#### 1.3.1 Data preprocessing

We implemented the data preprocessing stage of the protocol as a new R-package (*DecompPipeline*) that integrates quality filtering, adjustment for confounding factors and feature selection into a single, easy-to-use workflow. For loading, formatting and storing DNA methylation data we recommend our recently updated and extended *RnBeads* package.

##### Data import

Genome-wide DNA methylation can be profiled by different technologies such as whole-genome bisulfite sequencing (WGBS), reduced-representation bisulfite sequencing (RRBS) or the Illumina Infinium microarrays. In this protocol, we focus on microarray datasets due to their larger sample sizes that increase the efficiency of deconvolution. Nevertheless, our pipeline is similarly applicable to any other data type that provides DNA methylation calls at single CpG resolution. In addition to raw DNA methylation data, phenotypic information, such as age and sex, is required and converted into the internal *RnBeads*^26,27^ data structure. The input is checked for data quality using *RnBeads*’ reporting functionality.

##### Quality filtering

*DecompPipeline* performs quality-based filtering of CpGs across the samples in several steps (**Table 1**). First, CpGs are filtered according to a coverage threshold across the samples, and overall signal intensity (microarrays) or coverage (bisulfite sequencing) outliers are removed. Missing values can either be completely discarded from the dataset or imputed^37^. We further remove sites overlapping annotated or estimated single nucleotide polymorphisms (SNPs), sites on the sex chromosomes and cross-reactive sites^38^. Infinium data should be normalized prior to downstream analysis, and further sample properties can be inferred, such as the overall immune cell content using the LUMP algorithm^39^ or the epigenetic age^29^.

##### Covariate adjustment using ICA

DNA methylomes can be affected by various sources of variability, both of biological and technical nature that might mask the signals of interest. Deciding which of the detected components is of relevance for the investigated system and which components are associated with unwanted sources of variation (confounding factors) is a critical choice to be made by the user. For instance, components linked to age may be of interest in some investigated systems, while they may confound tumor heterogeneity in another setting. We use Independent Component Analysis (ICA)^30^ as a data-driven dimensionality reduction method that performs a matrix decomposition to adjust for a given set of confounders. ICA divides the experimentally observed data matrix *D*_*mn*_ into *k* independent signals *S*_*mk*_ mixed with the coefficients of *M*_*kn*_:

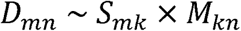

where *m* and *n* are the number of features (methylation sites) and samples, respectively. In contrast to the non-negative matrix factorization performed by *MeDeCom*, the entries of the matrices are not bounded to the [0, 1] interval. The weight matrix *M* can be linked to confounding factors, including sex, experimental batch effects or platform biases. As the method separates the original methylation profiles into statistically independent signals, the influence of these factors on the methylation profiles can be attributed to particular CpGs, which can either be removed or the weights (rows of *M*) of corresponding components can be set to zero^40^. Given that the influence of the investigated confounding factors is small, we recommend to set the corresponding components to zero and reconstruct an adjusted data matrix.

A pitfall of ICA that is specific to methylation data is the smoothing of the beta value distribution during ICA-based data transformation. Thus, a post-processing step is required to bring beta values into the expected range. To achieve this, reconstructed values are linearly rescaled in order to set the 1^st^ and 99^th^ percentiles to beta values of zero and one. Finally, in order to reduce stochasticity of ICA decomposition, we apply the consensus ICA approach^41^. ICA was run multiple times and the resulting matrices *S* and *M* were averaged between the runs. The stability of the components is estimated as the coefficient of determination (*R*^2^) between the columns of *S* observed in different runs.

##### Selection of informative CpG subsets

In order to obtain interpretable deconvolution results, further feature selection is required, since, for instance, lowly variable CpGs do not contribute to signature recovery, but add to the computational runtime. Using prior knowledge about the underlying cell types, for instance known cell type-specific CpGs, is the best option, given such knowledge is available^11,20^. In the absence of prior knowledge about the biological system of interest, typical strategies for feature selection include selecting the most highly variable sites, the ones with the highest loadings on the first few principal components, or a random selection. *DecompPipeline* provides 14 options to select CpG subsets (**Table 2**^42–44^), and multiple of these options can be included in a single execution of the pipeline.

**Table 2:** CpG selection options available in *DecompPipeline*.

| CPG SELECTION METHOD | CPG SUBSET SELECTED | DETAILS |
| --- | --- | --- |
| **VAR** | Most highly variable across the samples | *N_MARKERS* to determine the number of sites used |
| **RANDOM** | Random subset |  |
| **HYBRID** | Half as most highly variable, half randomly |  |
| **PCA** | Highest loadings on the first *N_PRIN_COMP* principal components | Default *N_PRIN_COMP=10* |
| **PCADAPT** | Principal Component Analysis implemented in the *bigstatsr* R-package | Privé *et al.*<sup>40</sup> |
| **ALL** | All that fulfill the quality criteria |  |
| **RANGE** | Largest range across the samples |  |
| **CUSTOM** | User-specified list |  |
| **ROWFSTAT** | Linked to given reference profiles using the F-statistics | Requires reference profiles |
| **PHENO** | Differentially methylated according to specified phenotypic groups using the *limma* method | Ritchie *et al.*<sup>38</sup> |
| **HOUSEMAN2012** | 50,000 listed as cell-type specific in the reference-based deconvolution method by Houseman *et al.* | Houseman *et al.*<sup>11</sup> |
| **HOUSEMAN2014** | According to the *RefFreeEWAS* method | Houseman *et al.*<sup>18</sup> |
| **JAFFE2014** | Cell-type specific in Jaffe *et al.* | Jaffe *et al.*<sup>39</sup> |
| **EDEC_STAGE0** | According to Stage 0 of the *EDec* approach | Requires reference profiles<br>Onuchic *et al.*<sup>19</sup> |

#### 1.3.2 Performing deconvolution using MeDeCom

Reference-free deconvolution methods, such as *RefFreeCellMix*^24^, *EDec*^20^ or *MeDeCom*^21^, estimate the matrix of latent methylation components (*T*) and the matrix of proportions (*A*) based on the DNA methylation matrix of sites selected in the previous step (*D*) by means of non-negative matrix factorization. We focus on *MeDeCom*, but the pipeline similarly supports *RefFreeCellMix* and *EDec. MeDeCom* optimizes the squared Frobenius norm of the difference between the true (measured) methylation matrix *D* and the matrix product of *T* and *A* (**Fig. 1**, middle). Desired properties of the factor matrices, i.e. restriction to the [0, 1] interval (*T* and *A*) with column sums equal to one (*A*), are enforced by respective constraints on their entries. Furthermore, *MeDeCom* penalizes the entries of *T* not equal to zero and one using quadratic regularization (maximal at entries equal to 0.5) controlled by the regularization parameter *λ*:

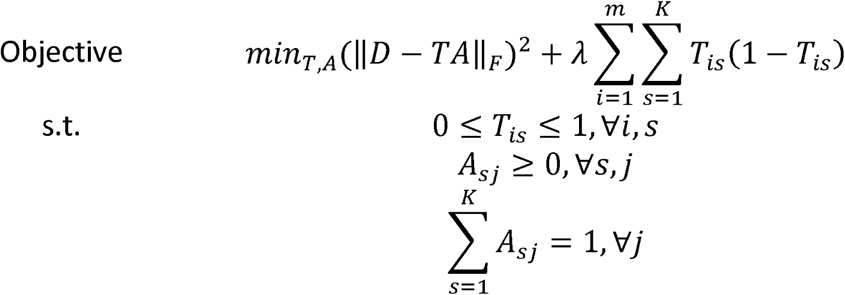

This optimization problem is solved using an alternating optimization scheme, which fixes either of *A* or *T* while fitting the other at each of its steps. Selecting suitable values for the regularization parameter (*λ*) and the number of latent components (*K*) is assisted by a cross-validation scheme that leaves out columns of *D*. Typically, a grid search for different values of *K* and *λ* is performed to determine the most suitable number of components and regularization, respectively. In order to reduce running time substantially, we recommend activating the parallel processing options on standalone workstations, or to use a high-performance computing cluster. The resulting solutions of the deconvolution problem are stored on disk and can be used for downstream interpretation.

#### 1.3.3 Interpretation of deconvolution results

In contrast to the reference-based case, interpretation of reference-free deconvolution results is non-trivial. *MeDeCom* produces a matrix of LMCs and a matched proportion matrix, both of which need to be biologically validated and interpreted. To facilitate an interactive interpretation, we created the semi-automated visualization tool *FactorViz. FactorViz* is an R/Shiny-based user interface with guidelines and functions for comprehensive biological inference. Initially, one of the possible *MeDeCom* solutions has to be chosen by selecting the parameters *K* and *λ* based on the cross-validation error. In order to investigate potential influences of covariates upon the estimated proportions and corresponding LMCs, the resulting proportion matrix is linked to technical or phenotypic traits using association tests. Furthermore, proportions can be linked to the expression of marker genes, or specific properties of the analyzed dataset such as survival time. To functionally annotate LMCs, we determine the sites that are specifically hypomethylated in a particular LMC in comparison to the median of the remaining LMCs, and treat the obtained sites as LMC-specific. Those sites are then used for GO^45^ and *LOLA*^46^ enrichment analysis in order to associate respective LMCs with functional categories, pathways and genomic features. Finally, the LMC matrix can be compared to available reference cell type profiles.

### 1.4 Level of expertise needed to implement the method

*DecompPipeline, MeDeCom*, and *FactorViz* are R-packages and thus require some minimal prior experience with the R programming language. Basic knowledge of the Unix command line interface is recommended for data handling. To follow the steps of this protocol, one only needs a few R function calls, but the function parameters need to be tailored to the target dataset. The graphical user interface *FactorViz* presents plots, which require a minimum knowledge of DNA methylation data and matrix factorization.

### 1.5 Limitations

Since *MeDeCom* tests the selected combinations of the regularization parameter *λ*, the number of LMCs *K* and several feature selection methods, the number of basic deconvolution jobs can reach 10,000 or more. Reference-free deconvolution is thus a computationally demanding task that requires high-performance computing infrastructure. When applied to larger datasets, the deconvolution can take several days to finish even on larger machines. Furthermore, the obtained LMC matrix needs to be biologically interpreted, which requires user interaction and input. A fully-automated biological inference of deconvolution results will be part of the next development steps of the protocol. Accounting for confounding factors, especially for those that might have a strong influence on the methylome, can lead to a substantially modified DNA methylation data matrix. The proposed pipeline provides diagnostic plots, but user interaction is still required to determine whether the effect of a particular covariate is to be removed.

## 2 Materials

### 2.1 Hardware

We recommend executing the proposed protocol on large systems with, e.g. 128 GB of main memory and 32 cores for a dataset of this size. For larger datasets, such as bisulfite sequencing data, we recommend to use a high-performance compute cluster or a cloud environment, such that the pipeline can automatically distribute jobs across different machines. Linux is the preferred operating system and the protocol was executed on a Debian Wheezy server with 32 cores and 128 GB of main memory.

### 2.2 Software

The proposed protocol requires a functional R installation, or Docker as an alternative. *MeDeCom, FactorViz* and *DecompPipeline* can be directly obtained from GitHub on Linux systems, and a binary release of *MeDeCom* has been deposited on GitHub for MacOS. See section *Data and Code Availability* below for URLs of the packages. Furthermore, to facilitate portability to other platforms, such as Windows 10, and in case of installation problems, we provide a Docker container (https://hub.docker.com/r/mscherer/medecom). See the instructions on the page of the Docker container on how to use the image. **Supplementary Table 2** lists operating systems and associated R-versions on which the protocol has been installed and tested.

### 2.3 Input data

We used publicly available data from The Cancer Genome Atlas (TCGA, https://www.cancer.gov/tcga) investigating lung adenocarcinoma (dataset TCGA-LUAD, https://portal.gdc.cancer.gov/legacy-archive/search/f) in 461 samples assayed using the Illumina 450k microarray, since lung cancer has high cellular and molecular heterogeneity^47^. The clinical metadata and the manifest file of the samples is available at https://portal.gdc.cancer.gov/projects/TCGA-LUAD and have been downloaded through the TCGA legacy archive on 2019-01-23. We used the Genomic Data Commons (GDC) data download tool (https://gdc.cancer.gov/access-data/gdc-data-transfer-tool) together with the manifest file to download the intensity data (IDAT) files and associated metadata.

## 3 Procedure

This protocol has been executed on a Linux system. For instructions on how to employ it on other operating systems, please see *Outline of the procedure and Materials*, and **Supplementary Table 2**.

### Installation TIMING 1 h

1. If R is not yet installed, follow the instructions at https://cran.r-project.org/. Create a working directory on a filesystem partition with sufficient storage capacity. Throughout the analysis, ∼20-30 Gb of free storage will be required. Be sure that all the files are downloaded into this working directory and that the code is executed within the directory.

2. Invoke R on the command line and install the *devtools* package. Install the software packages needed for deconvolution directly from GitHub: *MeDeCom, DecompPipeline* and *FactorViz*, and *RnBeads* from Bioconductor. ?TROUBLESHOOTING

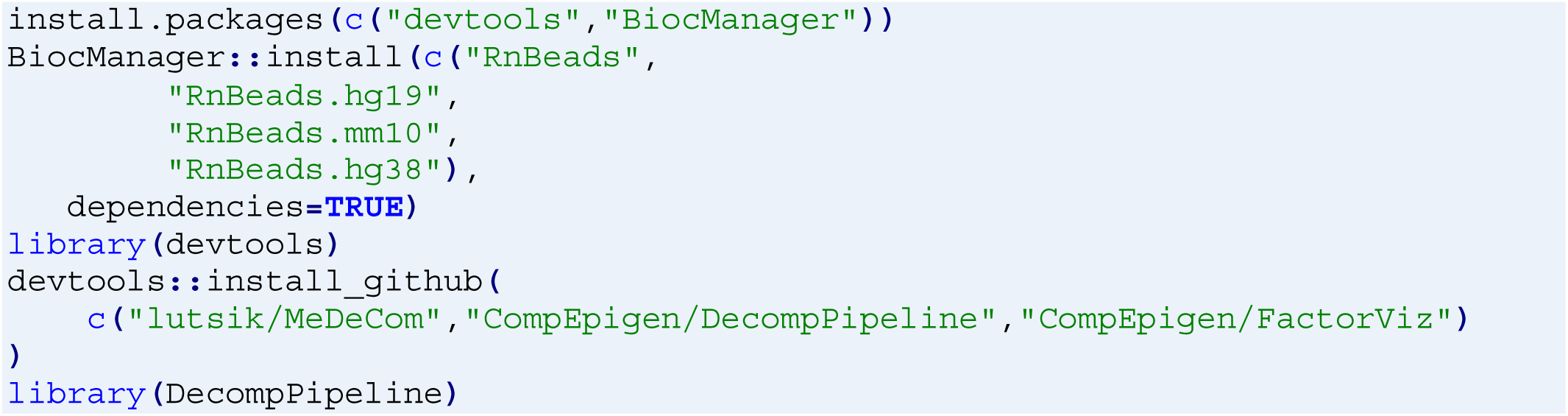

### Data retrieval TIMING 5 h

3. Use the Genomic Data Commons (GDC) data download tool (https://gdc.cancer.gov/access-data/gdc-data-transfer-tool) to download the IDAT files with the TCGA-LUAD manifest file (http://epigenomics.dkfz.de/DecompProtocol/data/gdc_manifest.2019-01-23.txt) and its associated metadata by running the download tool in a terminal session initiated in the working directory:

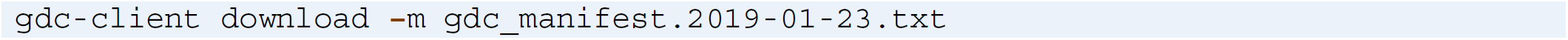

Store the IDAT in a new directory `idat`.

4. Retrieve the clinical metadata from https://portal.gdc.cancer.gov/projects/TCGA-LUAD by clicking on the *“Clinical”* button and unpacking the downloaded `.tar.gz` file into a new subdirectory `annotation` within your working directory. The remaining parsing is performed through an R session initiated in the same directory:

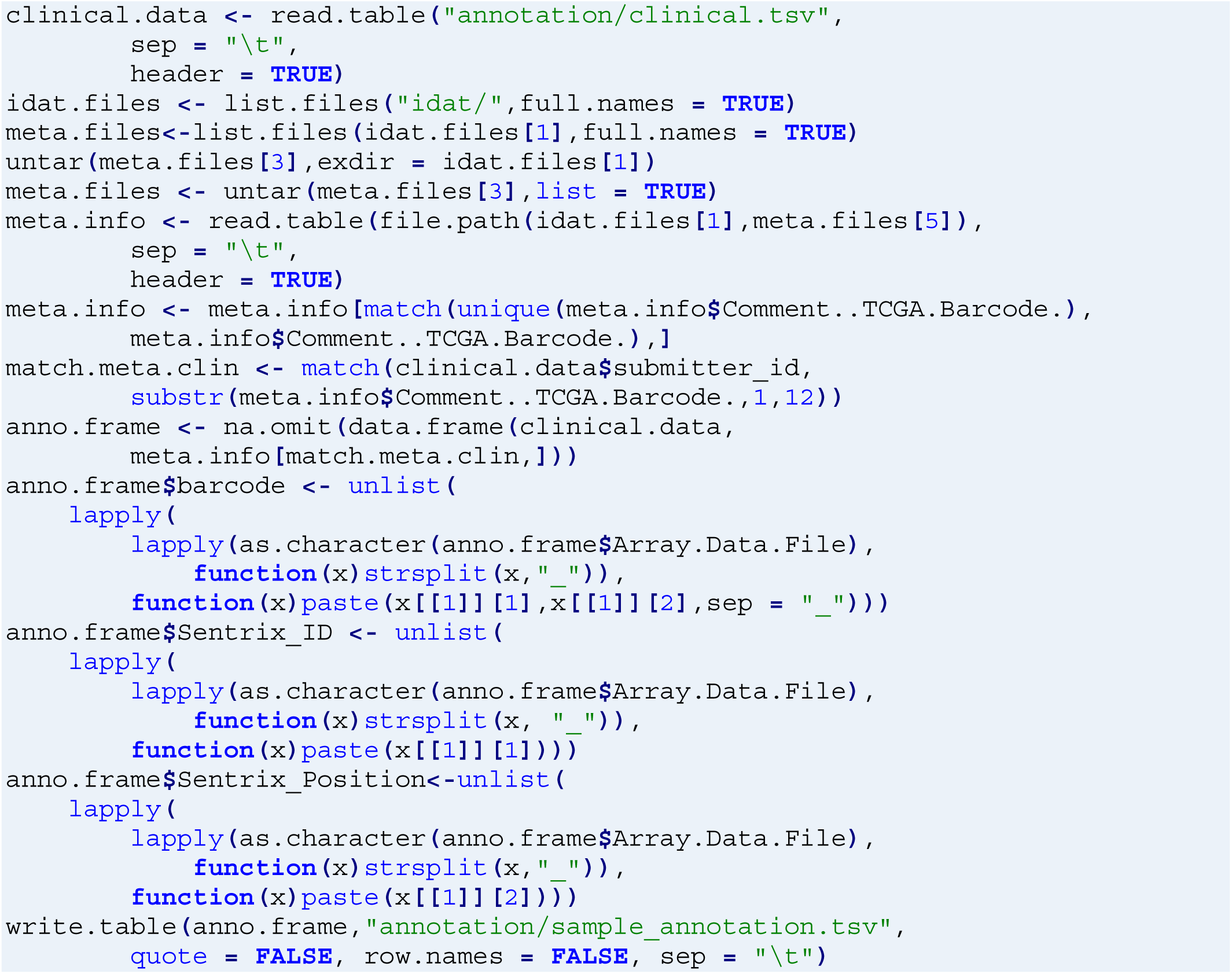

5. Copy the IDAT files into a single directory `idat` for downstream analysis.

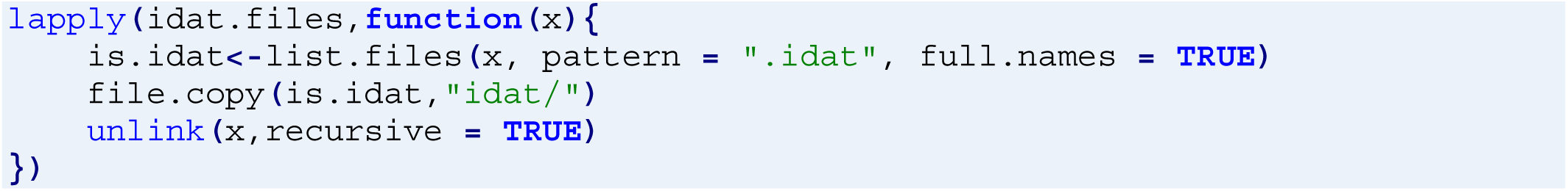

### Data import TIMING 2 h

6. *RnBeads* converts the files into a data object and performs basic quality control. Analysis options have to be specified for *RnBeads*, either through an XML file, or through the command line. Deactivate the preprocessing, exploratory, covariate inference, export and differential methylation modules, such that *RnBeads* only performs data import and quality control.

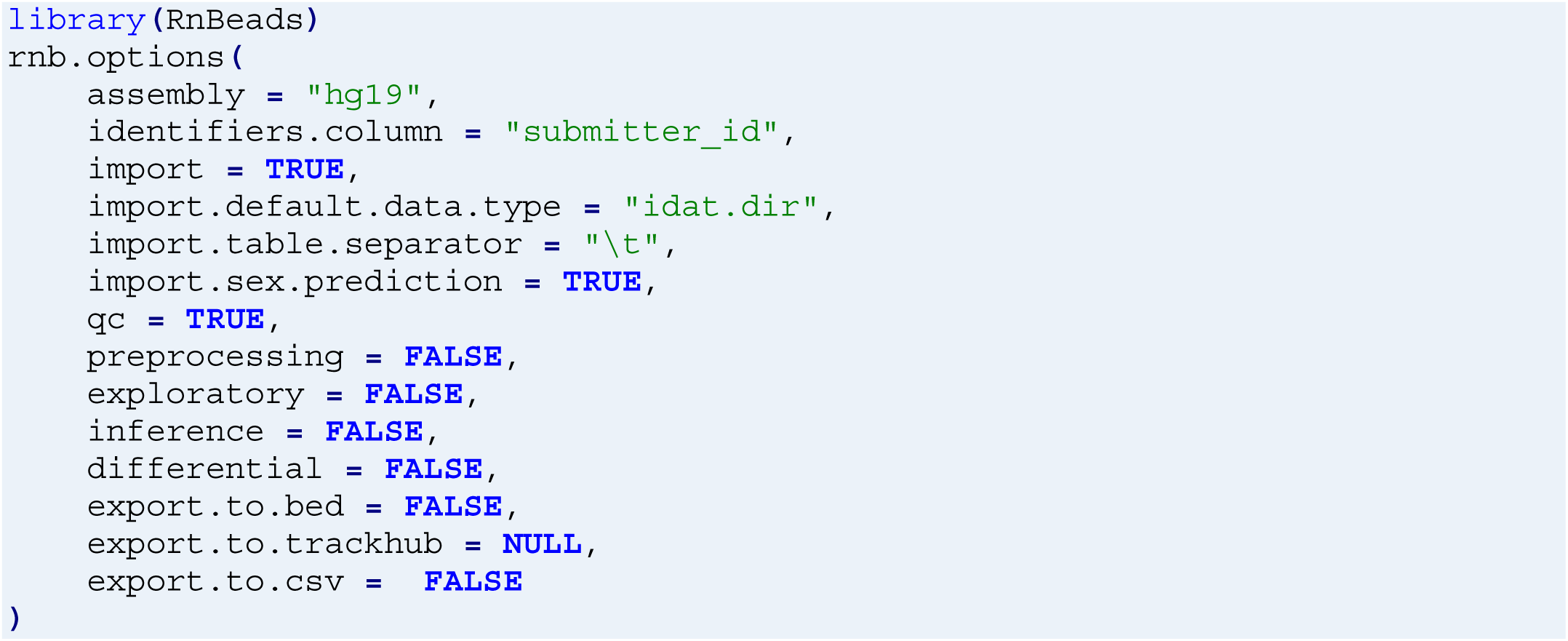

7. Specify the input to *RnBeads*: the sample annotation sheet created at the data retrieval step, the directory in which the IDAT files are stored and a directory to which the HTML report is to be saved. Additionally, specify a temporary directory and start the *RnBeads* analysis. ?TROUBLESHOOTING

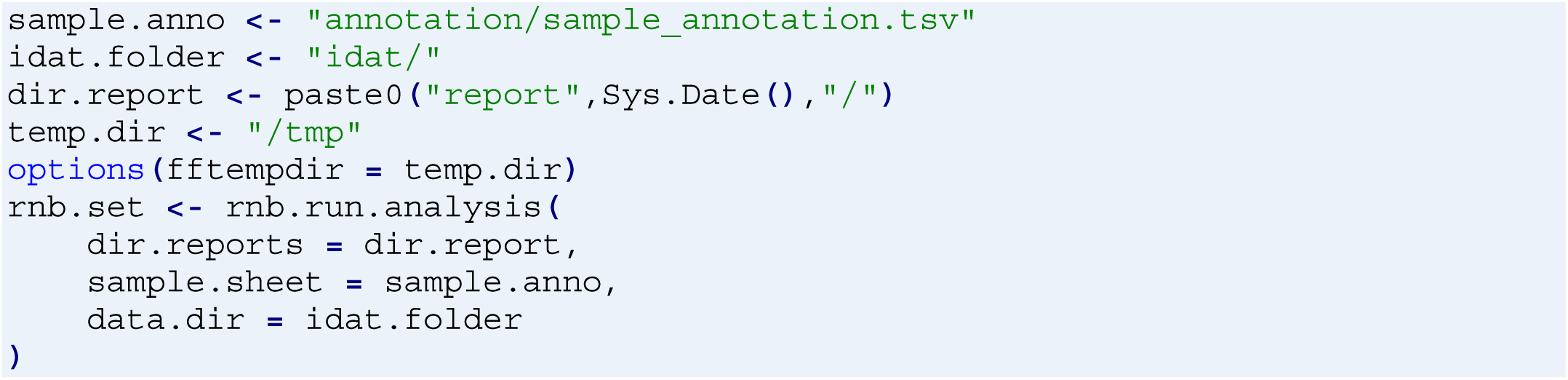

#### PAUSE POINT

The resulting object can be stored on disk and reloaded. We provide the intermediate result on our **Supplementary Website**: http://epigenomics.dkfz.de/downloads/DecompProtocol/rnbSet_unnormalized.zip.

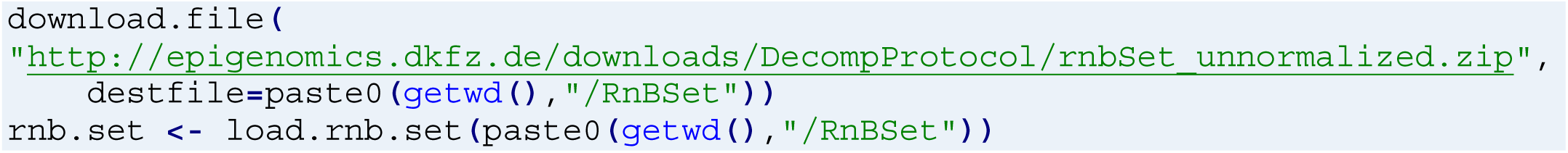

8. *RnBeads* creates an interactive HTML report, specifying the steps performed and the associated results. Data should meet the quality criteria in **Box 1** to be used for downstream analysis.

#### PAUSE POINT

For reference, we provide a complete RnBeads report on the supplementary website http://epigenomics.dkfz.de/downloads/DecompProtocol/RnBeads_Report_TCGA_LUAD/.

##### Box 1

###### Quality criteria for different quality control steps

**Control probes** All background control probes should have background intensity values in the range of 1,000-2,000. The low, medium and high control probes should show a substantially higher signal intensity in the range >10,000. Samples with high background intensity or low signal intensity should be discarded from the analysis.

**SNP methylation** The Illumina BeadArrays contain a few highly variable SNP probes, which should have methylation values close to 0, 0.5 and 1. In a genetically matched setup, samples from a similar genotype (e.g. the same family) should cluster together using the methylation values of these SNP probes. By this approach, potential sample mix-ups can be detected.

**Sex prediction** Patient sex can be reliably predicted from the signal intensities of the sites on the sex chromosomes. *RnBeads* trained a robust logistic regression classifier on a large training dataset. If predicted sex does not match the annotated sex, this indicates a sample mix-up.

### Preprocessing and filtering TIMING 22 h

9. For further data preparation and analysis steps use the *DecompPipeline* package. Processing options are provided through individual function parameters. Follow a stringent filtering strategy: (i) Filter CpGs covered by less than 3 beads, and probes that are in the 0.001 and 0.999 overall intensity quantiles (low and high intensity outliers). (ii) Remove all probes containing missing values in any of the samples. (iii) Discard sites outside of CpG context, overlapping annotated SNPs, located on the sex chromosomes and potentially cross-reactive probes. Finally, apply BMIQ normalization^48^ to account for the bias introduced by the two Infinium probe designs^49^. ?TROUBLESHOOTING

#### CRITICAL STEP

Removing too few or too many sites might have a strong influence on the final deconvolution results. Thus, we recommend to carefully check the available options (**Table 1**) and only change the default setting in case of low-quality data.

#### PAUSE POINT

We provide a list of selected CpGs as an intermediate result at http://epigenomics.dkfz.de/downloads/DecompProtocol/sites_passing_complete_filtering.csv.

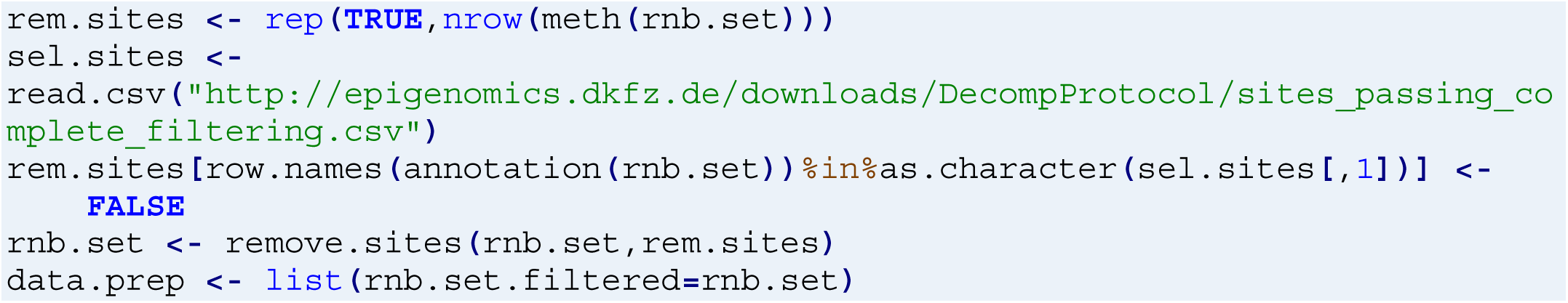

10. Adjustment for potential confounding factors is crucial in epigenomic studies and the influences of, for instance, donor sex, age, and genotype on the DNA methylation pattern are well-studied^28,29^. Use Independent Component Analysis (ICA, see Materials) to account for DNA methylation differences that are due to sex, age, race, and ethnicity. ?TROUBLESHOOTING

#### CRITICAL STEP

Confounding factor adjustment alters the overall data distribution, which might harm the overall bimodality of DNA methylation. Go to the analysis directory generated by *DecompPipeline* (`TCGA_LUAD`) and investigate the associations between Independent Components and the specified confounding factors. In case the reported p-value is lower than `alpha.fact`, ICA will automatically adjust for the corresponding confounding factor. Furthermore, inspect if the general bimodality of DNA methylation is preserved.

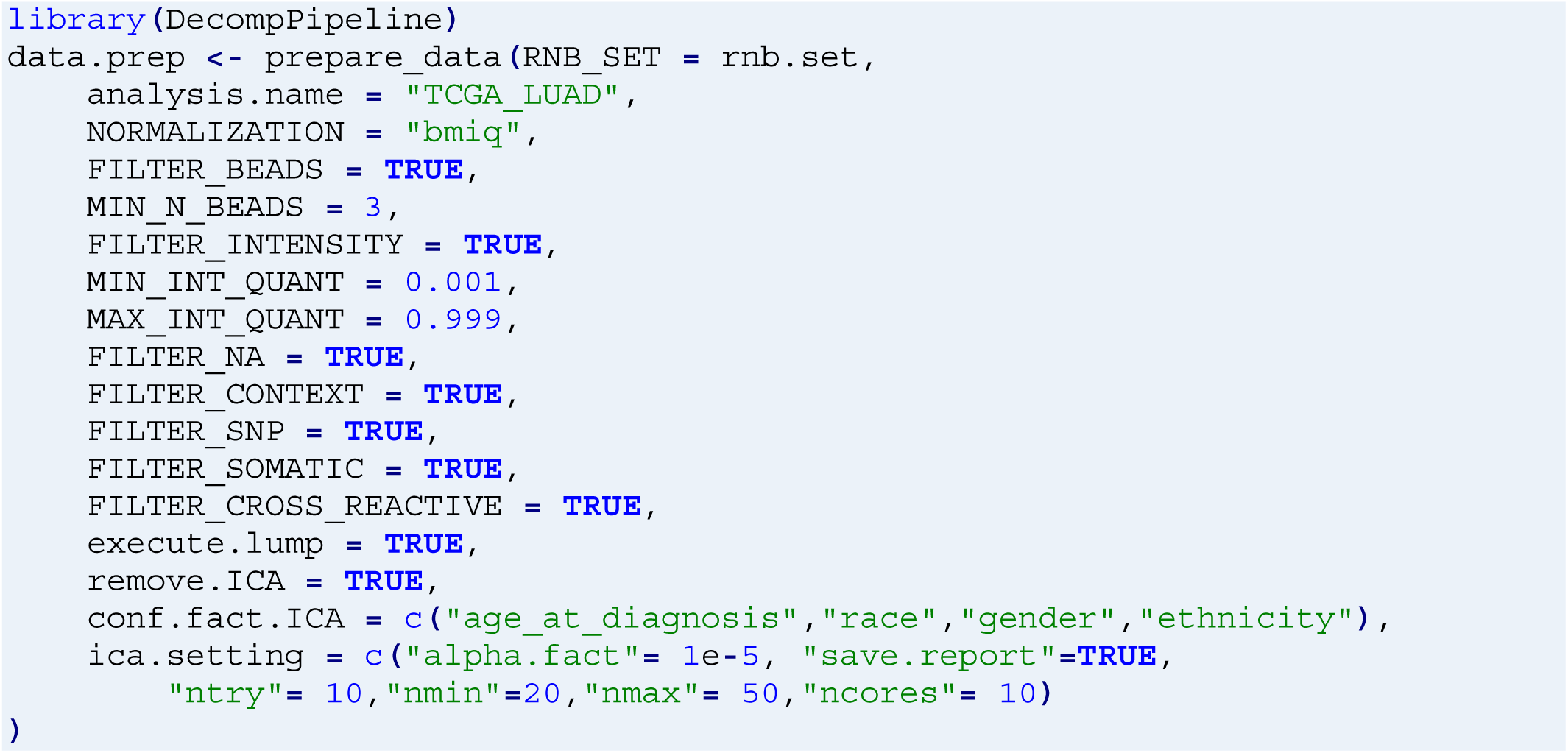

### Selection of CpG subsets TIMING 1 min

11. Select a subset of sites to be used for deconvolution. *DecompPipeline* provides different options (**Table 2**) through the *prepare_CG_subsets* function. Select the 5,000 most highly variable CpGs across the samples.

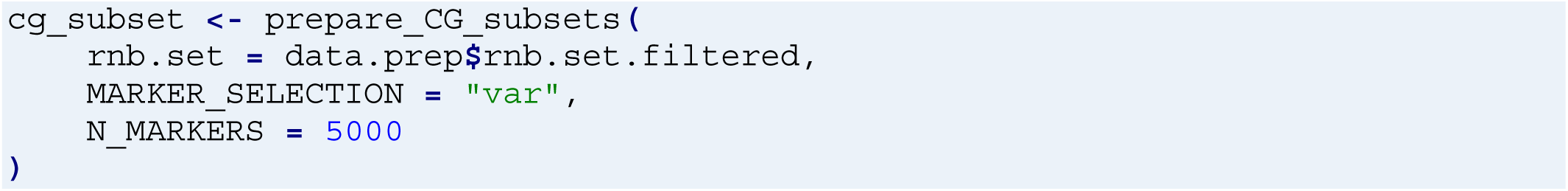

### Methylome deconvolution TIMING 54 h

12. Perform the deconvolution experiment. Use *MeDeCom* with a grid of values for the number of components (*K*) ranging from 2 to 15, which covers homogeneous to heterogeneous samples. Also, specify a grid for the regularization parameter (*λ*) from strong (0.01) to no regularization (0). ?TROUBLESHOOTING

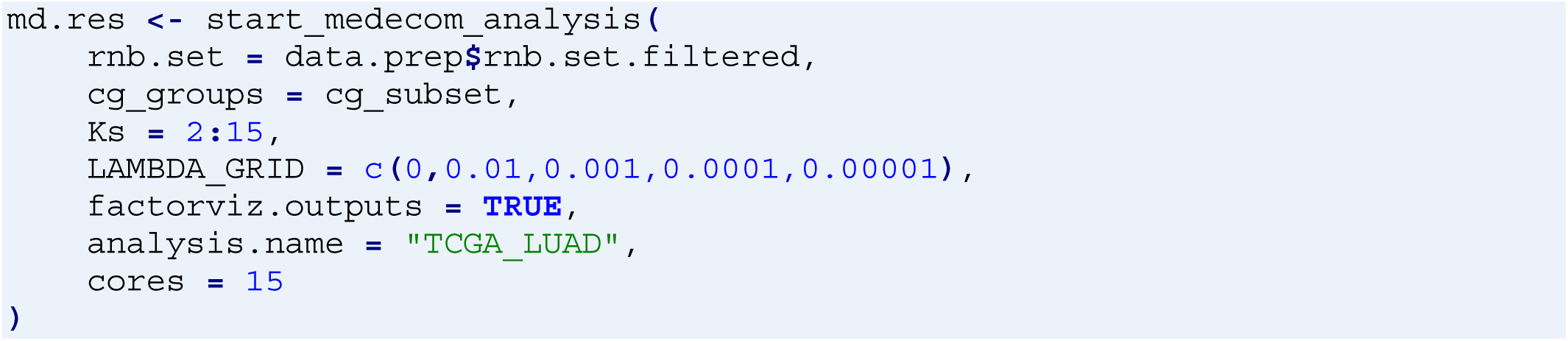

#### PAUSE POINT

The final *MeDeCom* result is stored in a format that can be directly imported with *FactorViz*. We provide the final result for exploration at http://epigenomics.dkfz.de/downloads/DecompProtocol/FactorViz_outputs.tar.gz.

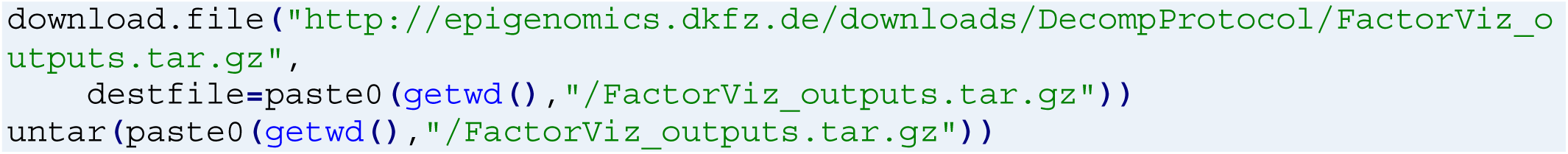

### Downstream analysis TIMING 3 h

13. Start the *FactorViz* application to visualize and interactively explore the deconvolution results (**Supplementary Fig. 1a, b**). ?TROUBLESHOOTING

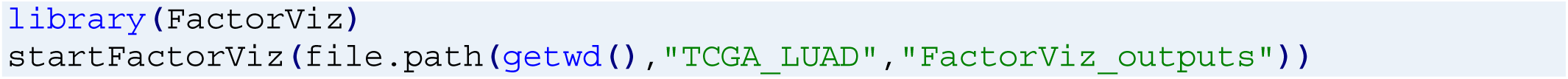

13a. In an additional R session started in the same working directory, deconvolution results can be loaded as an *MeDeComSet* object, which is part of the *FactorViz_outputs* directory, and can be used for additional analysis.

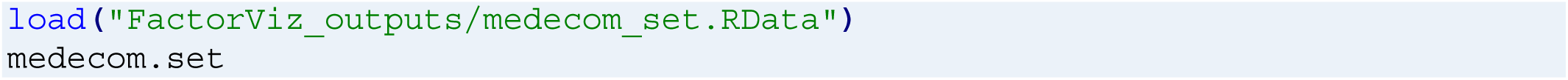

14. Determine the number of LMCs (K) and the regularization parameter (*λ*) based on the cross-validation error. (**Supplementary Fig. 1c, d**). First, go to panel “K selection” to plot cross-validation error for the range of *K*s specified earlier (**Supplementary Fig. 1c**). We recommend selecting *K* values as a trade-off between too high complexity, i.e. fitting the noise in the data (higher *K* values) and insufficient degrees of freedom (lower *K* values). This typically corresponds to the saddle point where cross-validation error starts to level out.

15. For a fixed *K*, select a value for the regularization parameter (*λ*) by proceeding to panel “Lambda selection” (**Supplementary Fig. 1d**). An optimal value for *λ* will often correspond to a local minimum of the cross-validation error. On the other hand, noticeable changes in other statistics, such as objective value or root mean squared error (RMSE), can point at a different *λ* value.

16. In the “Proportions” panel of *FactorViz*, visualize LMC proportions using heatmaps (**Supplementary Fig. 1e**). Proportion heatmaps can be visually annotated with available qualitative and quantitative traits (see dropdown “Color samples by”). Further visualization options include stacked barplot and lineplots.

17. In the “Meta-analysis” panel, associate LMC proportions with quantitative and qualitative traits using correlation- and t-tests (**Supplementary Fig. 1f**). Further sample annotations, such as mutational load (https://www.cbioportal.org/study/clinicalData?id=luad_tcga_pan_can_atlas_2018)^50^ and tumor purity scores can be obtained from public repositories and related publications^51^ (https://static-content.springer.com/esm/art%3A10.1038%2Fncomms3612/MediaObjects/41467_2013_BFncomms3612_MOESM489_ESM.xlsx). In addition, scripts to conduct such analysis are available on the **Supplementary Website** (http://epigenomics.dkfz.de/DecompProtocol/).

18. Explore the LMCs in the panel “LMCs” (**Supplementary Fig. 1g**). Several visualization options are available such as Multidimensional Scaling and histograms. Reference profiles can be used for joint visualization in case those are available.

19. Determine DNA methylation sites that are specifically hypo- and hypermethylated in an LMC by comparing the methylation values in the LMC matrix for each LMC to the median of the remaining LMCs in the “Meta-analysis” panel (**Supplementary Fig. 1h**). LMC-specific sites can be used for either a gene-centric Gene Ontology analysis (select “Enrichments” and “GO Enrichments” in dropdown “Analysis” and “Output type”) or for region-based enrichment analysis with the *LOLA* package (select “*LOLA* Enrichments” in dropdown “Output type”) (**Supplementary Fig. 1i, j**). The differential sites can be exported for further analysis, and the resulting plots can be stored by clicking on the “PDF” button.

#### PAUSE POINT

Additional built-in options for visualization of LMCs and proportions are available via the *MeDeCom* functions `plotLMCs` and `plotProportions` in R. Finally, the LMC and proportions matrix can be extracted from the *MeDeComSet object*, and the genomic annotation of CpGs (`ann.C`) can be obtained from the *FactorViz* output for a custom downstream analysis:

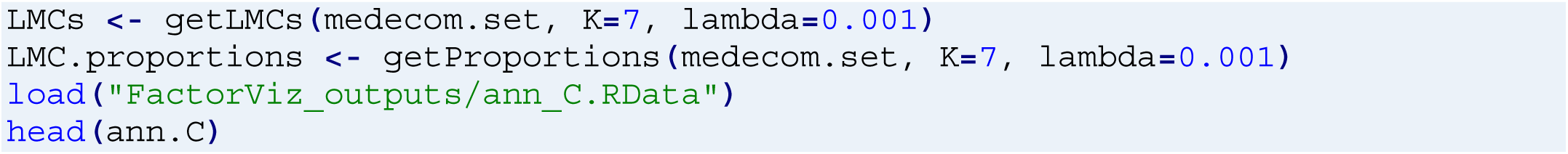

#### CRITICAL STEP

Interpretation of deconvolution results is crucial to obtain biological insights about the investigated system. In addition to the interpretation functions provided by *FactorViz*, prior knowledge such as marker gene expression values can be used to validate and interpret the deconvolution results. See a description of an example analysis comparing LMC proportions per sample to expression of epithelial, endothelial, stromal, and immune cell marker genes in matched RNA-seq data^52^ (**Supplementary Text**).

## 4 Troubleshooting

We provide troubleshooting advice in **Table 3**. For further support questions, feature requests or help, use GitHub’s issue system or write a mail to the tool developers.

**Table 3:**
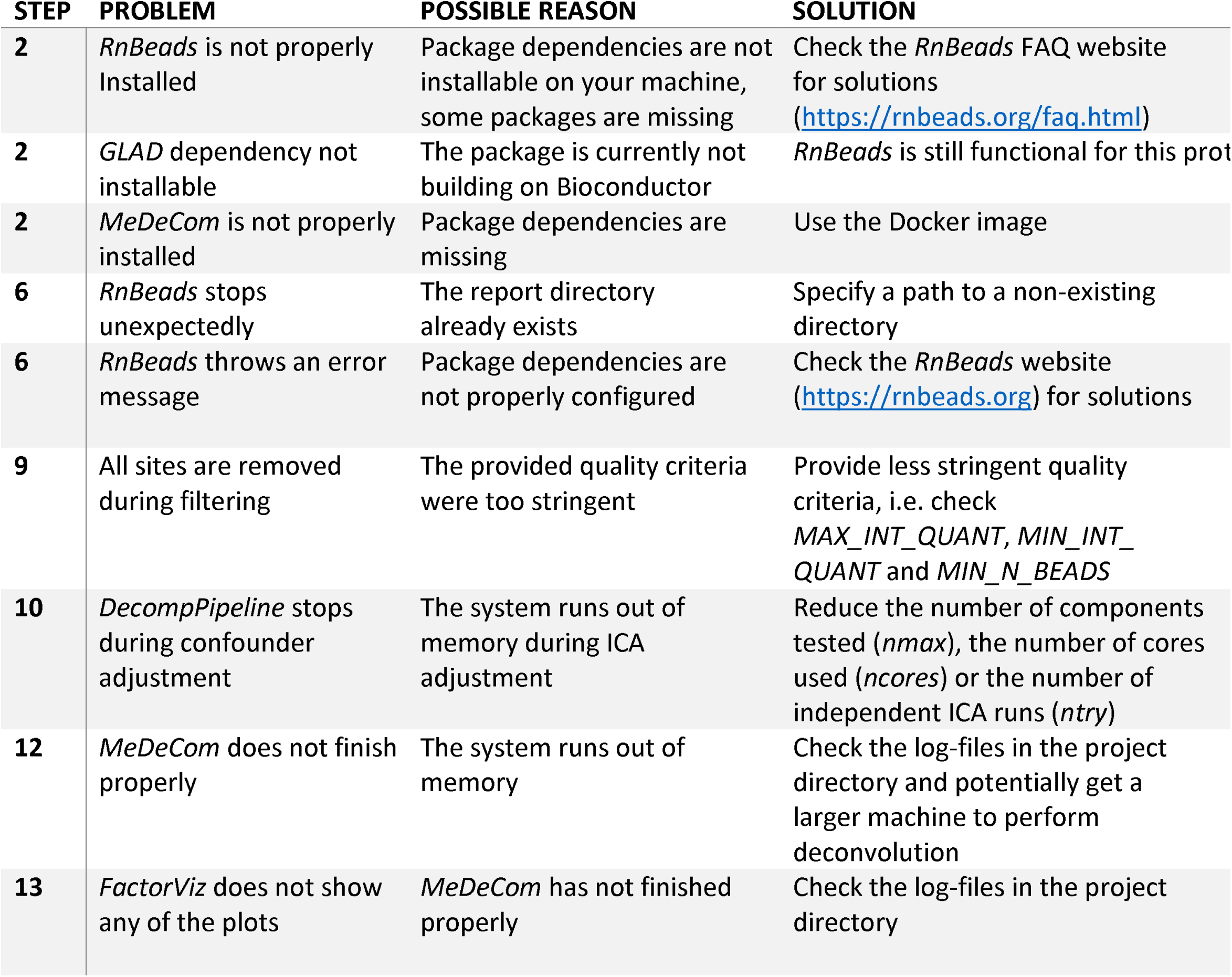
Troubleshooting for the individual steps of the reference-free deconvolution protocol.

## 5 Anticipated Results

### 5.1 Quality control and feature selection

Deconvolution analysis requires input DNA methylation data of sufficient technical quality, which we verified using *RnBeads*’ QC module. Quality control probes on the Infinium array did not reveal low-quality samples to be removed. Verifiable phenotypic information matched the inferred sample properties, such as predicted sex of all subjects (**Supplementary Fig. 2**). We used several criteria to select a set of high-confidence CpGs as input to *MeDeCom*. Most of the discarded sites (39.5 %) were removed, because they were covered by less than three beads in any of the samples, or showed unusually high or low intensity. Further CpG filtering steps, including sequence context (SNPs, sites on the sex chromosomes, 10.5 %) and removal of cross-reactive probes (2.5 %) eliminated further problematic sites. As a final outcome of the filtering procedure, 230,223 sites (47.4 % of 485,577) passed our stringent quality criteria and were used for downstream analysis.

**Fig. 2:**
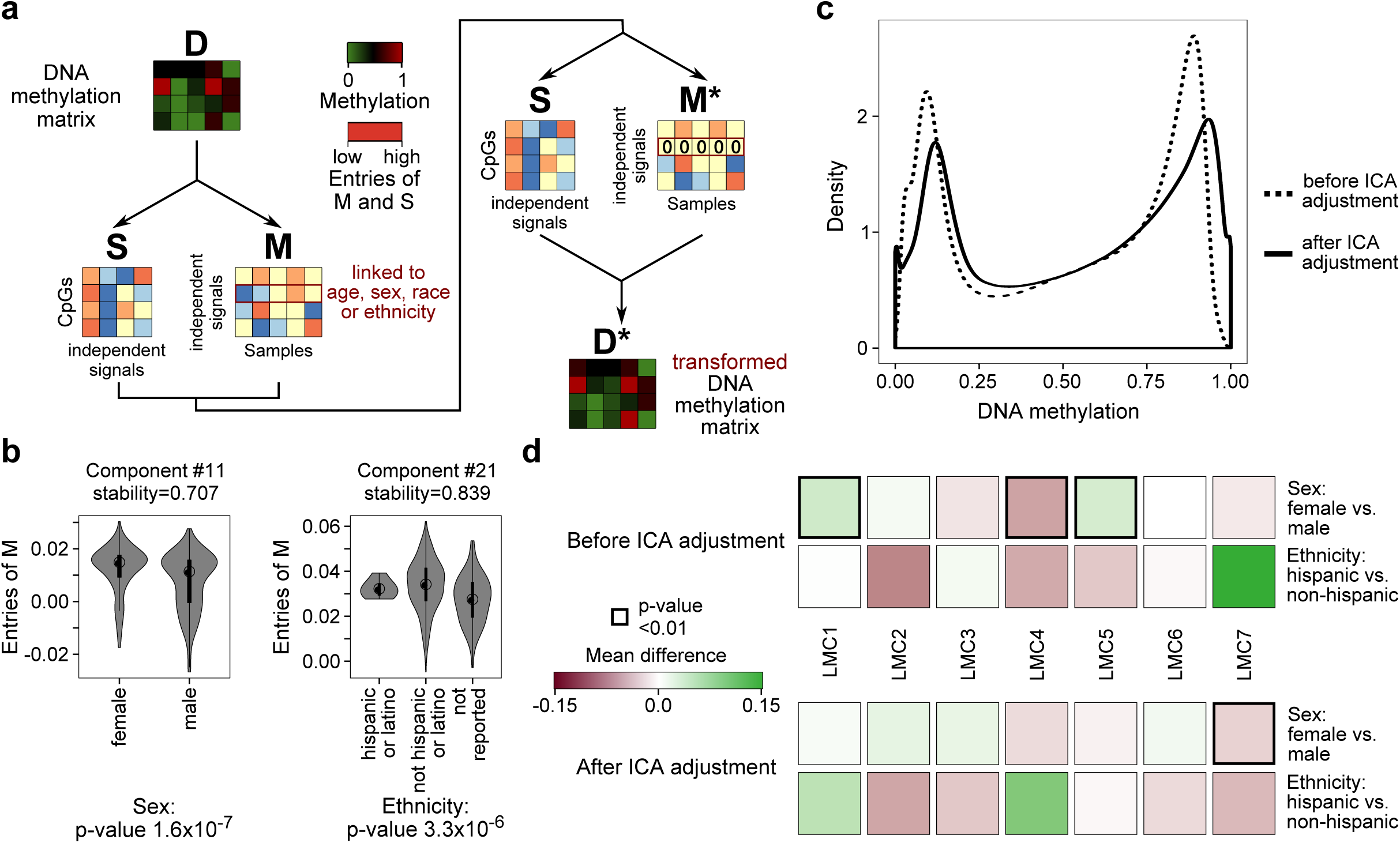
Evaluation of ICA on the TCGA LUAD dataset. **a** Overview of the ICA procedure. Components linked to confounding factors (here sex, age, ethnicity or race) are removed from the contribution matrix and an adjusted DNA methylation matrix is constructed. **b** Associations between the confounding factor sex and ethnicity with the entries of the proportion matrix *M* produced by ICA. P-values were computed using one-way ANOVA, points within the violin plots represent the median and the thick line 50 % of the samples. **c** Beta-value distributions of the transformed (*D*\*) and the untransformed (*D*) DNA methylation matrices. **d** Associations between LMC proportions and qualitative phenotypic traits. The color represents the absolute difference of the mean LMC proportions in the different groups defined by the phenotypic traits and significant p-values according to a two-sided t-test are indicated by a bold border.

### 5.2 Confounding factor analysis

We evaluated ICA by applying the proposed workflow to the TCGA dataset twice: once without adjusting for age, sex, race, and ethnicity, and once with the adjustment using ICA (**Fig. 2a**). ICA detected 22 components, of which two were significantly associated with sex and ethnicity, respectively (**Fig. 2b**). The overall distribution of the DNA methylation matrix was still bimodal after ICA adjustment, although there were notable peaks at methylation values zero and one, respectively. We argue that these peaks are a consequence of the scaling to the [0, 1] interval (**Fig. 2c**). After running *MeDeCom* independently on the unadjusted and the adjusted DNA methylation matrix, three of the detected components were significantly linked to sex in the unadjusted (p-values: LMC1: 6×10^−4^, LMC4: 1.4×10^−5^, LMC5: 3×10^−3^), but only one component was mildly linked to sex in the adjusted run (LMC7, p-value: 7.8×10^−4^, **Fig. 2d**). Although ICA component 11 was linked to ethnicity, we could not find a similar association with LMCs. Neither age nor race were significantly linked to any component produced by either ICA or *MeDeCom*.

### 5.3 Deconvolution results

The deconvolution results of the TCGA LUAD dataset are shown in **Fig. 3**. Since we did not have prior knowledge on the expected number of underlying cell types to select, we resorted to the cross-validation procedure of *MeDeCom*. We chose seven LMCs as the value of *K* at which the cross-validation error started to level out (**Supplementary Fig. 3a**). Similarly, we selected λ=0.001 (regularization parameter) as the point where cross-validation error is still low, while objective value and RMSE substantially change (**Supplementary Fig. 3b**). Notably, LMC5 was particularly hypomethylated and LMC6 showed a high overall methylation level, while the remaining LMCs were rather intermediately methylated (**Supplementary Fig. 3c, d**).

**Fig. 3:**
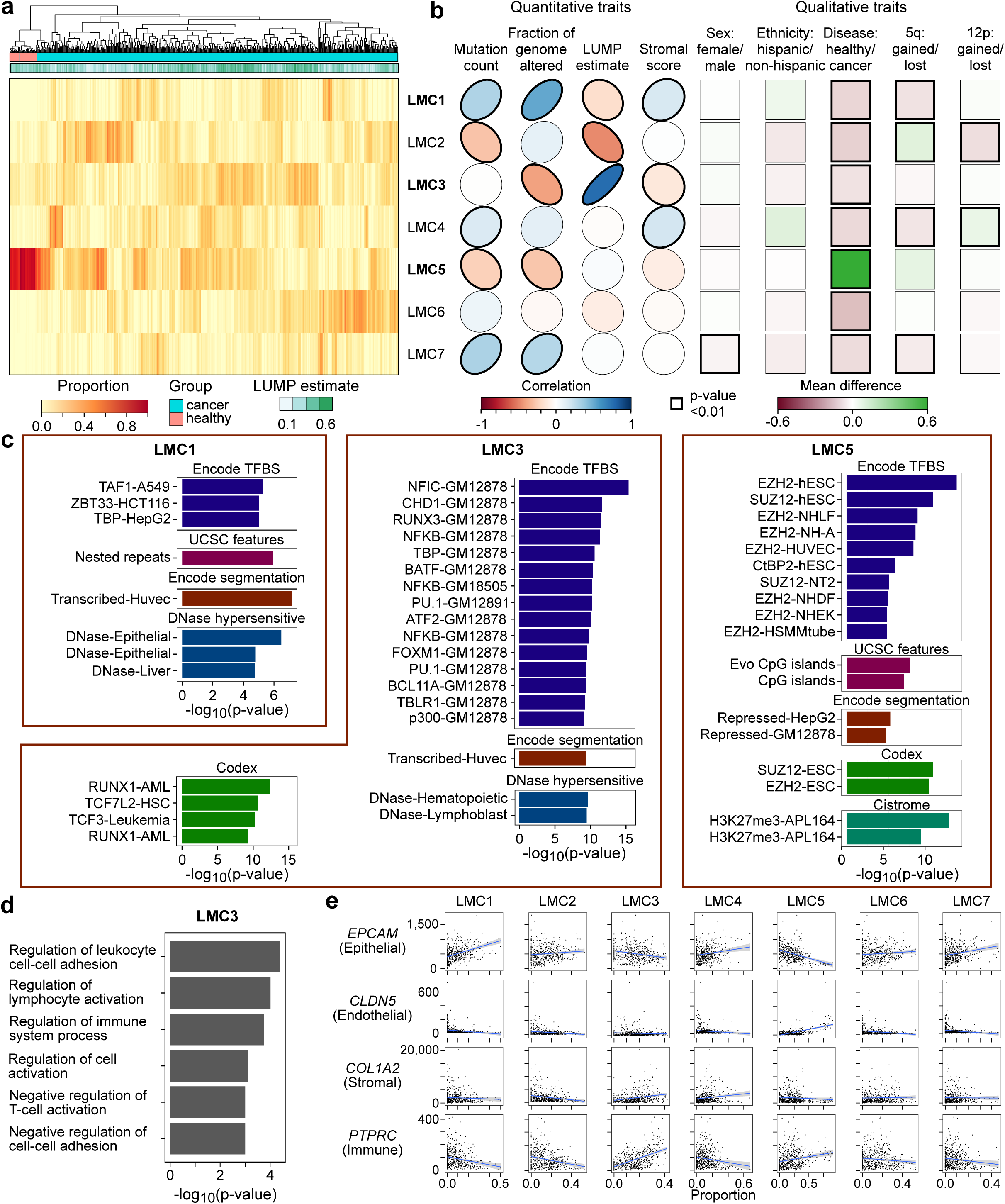
Interpreting *MeDeCom* results with *FactorViz*. **a** Proportion heatmap of LMCs in the different samples. We selected K=7 LMCs and set λ=0.001. The samples were hierarchically clustered according to the Euclidean distance between the proportions using complete linkage. We annotated samples using disease status and with the sample-specific LUMP estimate. **b** Associations between the phenotypic traits and LMC proportions. For quantitative traits, the Pearson correlations are shown as ellipses that are directed to the upper right for positive and to the lower right for negative correlations, respectively. For qualitative traits, the absolute difference of the proportions in the two groups (e.g. female vs. male) is shown. P-values (two-sided correlation test for quantitative and two-sided t-test for categorical variables) less than 0.01 are indicated by bold borders. *LOLA*^*46*^ (**c**) and *GO*^*45*^ (**d**) enrichment analysis of the LMC-specific hypomethylated sites for LMCs 1, 3 and 5. No significant GO enrichment was found for LMC 1 and 5. Sites were defined as LMC-specific hypomethylated if the difference between the LMC value and the median of all other LMCs was lower than 0.5. P-values have been adjusted for multiple testing with the Benjamini-Hochberg method^58^. **e** Scatterplots between LMC proportions per sample and known marker gene expression of different lung cell types. The gene expression was measured using counts per million (CPM).

We further investigated the biological implications of the detected LMCs. To this end, we found that LMC5 had substantially higher proportions in the healthy tissue samples (**Fig. 3a**). This indicated that reference-free deconvolution was able to capture the inherent methylation signatures specific to cancerous and healthy tissues. When we conducted enrichment analysis for the sites that were specifically hypomethylated in LMC5, we found that transcription factor binding sites for the Polycomb repressive complex (*SUZ12, EZH2*) were overrepresented. Polycomb repressive complex, which typically represses oncogenes, is often dysregulated in cancers; a process that is often associated with hypermethylation^53–55^. What is more, proportions of LMC5 in tumor tissues provided a generic estimate of tumor sample purity, i.e. degree of contamination by adjacent normal tissue. Thus, we were able to capture tumor-specific methylation signatures without conducting differential analysis between two phenotypic groups.

LMC3 showed highly variable proportions across the samples and was the main driver of the overall cancer sample clustering. LMC3 proportions were strongly correlated with the LUMP estimate (**Fig. 3b**), which predicts the overall immune cell content of a sample^39^. Furthermore, we detected enrichments of LMC3-specific hypomethylated sites towards leukocyte (B-lymphocyte) specific transcription factor binding sites and immune response related GO terms (**Fig. 3c, d, Supplementary Fig. 4**). We concluded that LMC3 most likely represented tumor infiltrating immune cells. The extent of tumor infiltration might be relevant to associate cancer state to patient prognosis^56^.

To determine whether the detected LMCs reflected the expression of known marker genes of lung tissue cell types, we selected *EPCAM* as an epithelial, *CLDN5* as an endothelial, *COL1A2* as a stromal, and *PTPRC* as an immune cell marker^52^. We collected gene expression data from TCGA for the samples (data processing is described in the **Supplementary Text**) and found LMC3 to be correlated to *PTPRC* expression. Furthermore, LMC1 was strongly associated with the epithelial marker expression and LMC5 with the endothelial marker *CLDN5* (**Fig. 3e**).

Many of the detected components were linked to cancer mutational status, such as overall mutation count or chromosomal gain or loss (**Supplementary Fig. 5**), or cancer stemness (**Supplementary Fig. 6**). For instance, LMC1 was inversely linked to copy number gain of chromosome 1p and 16p. We further investigated the sites that were particularly hypo- or hypermethylated in LMC4, since LMC4 showed a high proportion in a small subset of samples that clustered separately from the remaining cancer samples (**Fig. 3a**). One of these sites, cg26992600, is located 3 kb upstream of the TSS of NKX2-8, a gene that has potential roles in the progression of lung cancers^57^ (**Supplementary Fig. 7, Supplementary Table 3**). Finally, in a survival analysis of LMC proportions, we found that proportions of LMC5 and LMC6 were associated with survival time (p-values: LMC5: 0.18, LMC6: 0.03, **Supplementary Fig. 8**), warranting further investigation of their biological nature.

## Conclusions

Taken together, our methylome deconvolution protocol was able to provide novel biological insights about the cellular and intra-tumor heterogeneity of lung cancer. We expect the protocol to be of great benefit to all investigators performing DNA methylation analysis in complex and underexplored experimental systems, including bulk samples of highly heterogeneous tissues and tumors.

## Acknowledgements

We thank the HADACA consortium (Health Data Challenge, Aussois, Dec 2018) for valuable input and Divanshu Gupta for thoroughly testing the proposed pipeline. We are grateful to Kersten Breuer for testing the Docker container, and to Francisco Azuaje for supporting the collaboration.

## Author contributions

M.S. and P.L. implemented most of the computational procedures. P.L. previously developed and published *MeDeCom*. S.S, M.S., and P.L. implemented *FactorViz*. P.N. and T.K. implemented consensus ICA. M.S. performed the analysis of the example datasets and created all figures and tables. P.N., R.T. and V.M. provided crucial input to the analysis and interpretation, and thoroughly tested the protocol. P.L., J.W., T.L., and C.P. jointly supervised the project. M.S. and P.L. wrote the manuscript, with contributions of all co-authors. All authors read and approved the final text.

## Data and code availability

The results shown here are wholly or partially based upon data generated by the TCGA (TCGA-LUAD dataset) Research Network: https://www.cancer.gov/tcga.

All R packages are available from public code repositories:

*DecompPipeline*: https://github.com/CompEpigen/DecompPipeline

*MeDeCom:* https://github.com/lutsik/MeDeCom

*FactorViz*: *https://github.com/CompEpigen/FactorViz*

*consensusICA*: https://gitlab.com/biomodlih/consica.

The pipeline behind our protocol is also implemented as a Docker container available from *DockerHub*: https://hub.docker.com/r/mscherer/medecom. Supplementary resources and R scripts used to generate the figures are available from http://epigenomics.dkfz.de/DecompProtocol/.

## Supplementary Material

**Supplementary File 1**: PDF comprising Supplementary Tables, Figures and Text.

**Supplementary Website:** Additional resources for the protocol, intermediate results and useful R code snippets are available at http://epigenomics.dkfz.de/DecompProtocol/.

## References

1. Durek, P. et al. Epigenomic Profiling of Human CD4+ T Cells Supports a Linear Differentiation Model and Highlights Molecular Regulators of Memory Development. Immunity 45, 1148–1161 (2016).

2. Karpinski, P., Pesz, K. & Sasiadek, M. M. Pan-cancer analysis reveals presence of pronounced DNA methylation drift in CpG island methylator phenotype clusters. Epigenomics 9, 1341–1352 (2017).

3. Møller, M. et al. Heterogeneous patterns of DNA methylation-based field effects in histologically normal prostate tissue from cancer patients. Sci. Rep. 7, 40636 (2017).

4. Vidal, E. et al. A DNA methylation map of human cancer at single base-pair resolution. Oncogene 36, 5648–5657 (2017).

5. Azuara, D. et al. New Methylation Biomarker Panel for Early Diagnosis of Dysplasia or Cancer in High-Risk Inflammatory Bowel Disease Patients. Inflamm. Bowel Dis. 24, 2555–2564 (2018).

6. Horvath, S. & Raj, K. DNA methylation-based biomarkers and the epigenetic clock theory of ageing. Nat. Rev. Genet. 19, 371–384 (2018).

7. Stunnenberg, H. G. et al. The International Human Epigenome Consortium: A Blueprint for Scientific Collaboration and Discovery. Cell 167, 1145–1149 (2016).

8. Adams, D. et al. BLUEPRINT to decode the epigenetic signature written in blood. Nat. Biotechnol. 30, 224–226 (2012).

9. Bock, C. Analysing and interpreting DNA methylation data. Nat. Rev. Genet. 13, 705–719 (2012).

10. Teschendorff, A. E. & Relton, C. L. Statistical and integrative system-level analysis of DNA methylation data. Nat. Rev. Genet. 19, 129–147 (2017).

11. Houseman, E. A. et al. DNA methylation arrays as surrogate measures of cell mixture distribution. BMC Bioinformatics 13, 86 (2012).

12. Teschendorff, A. E., Breeze, C. E., Zheng, S. C. & Beck, S. A comparison of reference-based algorithms for correcting cell-type heterogeneity in Epigenome-Wide Association Studies. BMC Bioinformatics 18, 105 (2017).

13. Chakravarthy, A. et al. Pan-cancer deconvolution of tumour composition using DNA methylation. Nat. Commun. 9, 3220 (2018).

14. Hicks, S. C. & Irizarry, R. A. methylCC: technology-independent estimation of cell type composition using differentially methylated regions. Genome Biol. 20, 261 (2019).

15. Kaushal, A. et al. Comparison of different cell type correction methods for genome-scale epigenetics studies. BMC Bioinformatics 18, 216 (2017).

16. Zou, J., Lippert, C., Heckerman, D., Aryee, M. & Listgarten, J. Epigenome-wide association studies without the need for cell-type composition. Nat. Methods 11, 309–311 (2014).

17. Rahmani, E. et al. Sparse PCA corrects for cell type heterogeneity in epigenome-wide association studies. Nat. Methods 13, 443–445 (2016).

18. Rahmani, E. et al. BayesCCE: a Bayesian framework for estimating cell-type composition from DNA methylation without the need for methylation reference. Genome Biol. 19, 141 (2018).

19. Houseman, E. A., Molitor, J. & Marsit, C. J. Reference-free cell mixture adjustments in analysis of DNA methylation data. Bioinformatics 30, 1431–1439 (2014).

20. Onuchic, V. et al. Epigenomic Deconvolution of Breast Tumors Reveals Metabolic Coupling between Constituent Cell Types. Cell Rep. 17, 2075–2086 (2016).

21. Lutsik, P. et al. MeDeCom: discovery and quantification of latent components of heterogeneous methylomes. Genome Biol. 18, 55 (2017).

22. Rahmani, E. et al. Cell-type-specific resolution epigenetics without the need for cell sorting or single-cell biology. Nat. Commun. 10, 3417 (2019).

23. Thompson, M., Chen, Z. J., Rahmani, E. & Halperin, E. CONFINED: Distinguishing biological from technical sources of variation by leveraging multiple methylation datasets. Genome Biol. 20, 138 (2019).

24. Houseman, E. A. et al. Reference-free deconvolution of DNA methylation data and mediation by cell composition effects. BMC Bioinformatics 17, 259 (2016).

25. Decamps, C. et al. Guidelines for cell-type heterogeneity quantification based on a comparative analysis of reference-free DNA methylation deconvolution software. Preprint at https://www.biorxiv.org/content/10.1101/698050v1 (2019).

26. Assenov, Y. et al. Comprehensive analysis of DNA methylation data with RnBeads. Nat. Methods 11, 1138–1140 (2014).

27. Müller, F. et al. RnBeads 2.0: comprehensive analysis of DNA methylation data. Genome Biol. 20, 55 (2019).

28. Heyn, H. et al. Distinct DNA methylomes of newborns and centenarians. Proc. Natl. Acad. Sci. 109, 10522–10527 (2012).

29. Horvath, S. DNA methylation age of human tissues and cell types. Genome Biol. 14, R115 (2013).

30. Sompairac, N. et al. Independent Component Analysis for Unraveling the Complexity of Cancer Omics Datasets. Int. J. Mol. Sci. 20, 4414 (2019).

31. Everson, T. M. et al. Cadmium-associated differential methylation throughout the placental genome: Epigenome-wide association study of two U.S. birth cohorts. Environ. Health Perspect. 126, 1–13 (2018).

32. Carlström, K. E. et al. Therapeutic efficacy of dimethyl fumarate in relapsing-remitting multiple sclerosis associates with ROS pathway in monocytes. Nat. Commun. 10, 3081 (2019).

33. Goeppert, B. et al. Integrative Analysis Defines Distinct Prognostic Subgroups of Intrahepatic Cholangiocarcinoma. Hepatology 69, 2091–2106 (2019).

34. Man, Y. G. et al. Tumor-infiltrating immune cells promoting tumor invasion and metastasis: Existing theories. J. Cancer 4, 84–95 (2013).

35. Luo, C. et al. Single-cell methylomes identify neuronal subtypes and regulatory elements in mammalian cortex. Science (80-.). 357, 600–604 (2017).

36. Mulqueen, R. M. et al. Highly scalable generation of DNA methylation profiles in single cells. Nat. Biotechnol. 36, 428–431 (2018).

37. Troyanskaya, O. et al. Missing value estimation methods for DNA microarrays. Bioinformatics 17, 520–525 (2001).

38. Chen, Y. A. et al. Discovery of cross-reactive probes and polymorphic CpGs in the Illumina Infinium HumanMethylation450 microarray. Epigenetics 8, 203–209 (2013).

39. Aran, D., Sirota, M. & Butte, A. J. Systematic pan-cancer analysis of tumour purity. Nat. Commun. 6, 8971 (2015).

40. Dirkse, A. et al. Stem cell-associated heterogeneity in Glioblastoma results from intrinsic tumor plasticity shaped by the microenvironment. Nat. Commun. 10, 1787 (2019).

41. Nazarov, P. V et al. Deconvolution of transcriptomes and miRNomes by independent component analysis provides insights into biological processes and clinical outcomes of melanoma patients. BMC Med. Genomics 12, 132 (2019).

42. Ritchie, M. E. et al. limma powers differential expression analyses for RNA-sequencing and microarray studies. Nucleic Acids Res. 43, e47 (2015).

43. Jaffe, A. E. & Irizarry, R. A. Accounting for cellular heterogeneity is critical in epigenome-wide association studies. Genome Biol. 15, R31 (2014).

44. Prive, F., Aschard, H., Ziyatdinov, A. & Blum, M. G. B. Efficient analysis of large-scale genome-wide data with two R packages: Bigstatsr and bigsnpr. Bioinformatics 34, 2781–2787 (2018).

45. Falcon, S. & Gentleman, R. Using GOstats to test gene lists for GO term association. Bioinformatics 23, 257–258 (2007).

46. Sheffield, N. & Bock, C. LOLA:Enrichment analysis for genomic region sets and regulatory elements in R and Bioconductor. Bioinformatics 32, 587–589 (2016).

47. Testa, U., Castelli, G. & Pelosi, E. Lung cancers: Molecular characterization, clonal heterogeneity and evolution, and cancer stem cells. Cancers (Basel). 10, 1–81 (2018).

48. Teschendorff, A. E. et al. A beta-mixture quantile normalization method for correcting probe design bias in Illumina Infinium 450 k DNA methylation data. Bioinformatics 29, 189–196 (2013).

49. Bibikova, M. et al. High density DNA methylation array with single CpG site resolution. Genomics 98, 288–295 (2011).

50. Cerami, E. et al. The cBio Cancer Genomics Portal: An open platform for exploring multidimensional cancer genomics data. Cancer Discov. 2, 401–404 (2012).

51. Yoshihara, K. et al. Inferring tumour purity and stromal and immune cell admixture from expression data. Nat. Commun. 4, 2612 (2013).

52. Sit, R. V, Chang, S., Conley, S. D., Mori, Y. & Seita, J. A molecular cell atlas of the human lung from single cell RNA sequencing. Preprint at https://www.biorxiv.org/content/10.1101/742320v1 (2019)

53. Hahn, M. A. et al. Methylation of Polycomb target genes in intestinal cancer is mediated by inflammation. Cancer Res. 68, 10280 (2008).

54. Varambally, S. et al. The polycomb group protein EZH2 is involved in progression of prostate cancer. Nature 419, 624–629 (2002).

55. Cai, Y. et al. Epigenetic alterations to Polycomb targets precede malignant transition in a mouse model of breast cancer. Sci. Rep. 8, 1–12 (2018).

56. Ward, M. J. et al. Tumour-infiltrating lymphocytes predict for outcome in HPV-positive oropharyngeal cancer. Br. J. Cancer 110, 489–500 (2014).

57. Harris, T. et al. Both gene amplification and allelic loss occur at 14q13.3 in lung cancer. Clin. Cancer Res. 17, 690–699 (2011).

58. Benajmini, Y. & Hochberg, Y. Controlling the False Discovery Rate: A Practical and Powerful Approach to Multiple Testing. J. R. Stat. Soc. Ser. B 57, 289–300 (1995).

