## Supplementary Data for "Reference-free deconvolution of complex DNA methylation data – a systematic protocol"

### Supplementary Material To
### Deconvolution of complex DNA methylation data - a detailed protocol

Michael Scherer, Petr V. Nazarov, Reka Toth, Shashwat Sahay, Tony Kaoma, Valentin Maurer, Christoph Plass, Thomas Lengauer, Jörn Walter, and Pavlo Lutsik

February 14, 2020

#### Supplementary Text

##### Gene expression data processing

1. Download matched RNA-seq data from the TCGA legacy archive using the *TCGAbiolinks* [1] R package as normalized results.

```
library(TCGAbiolinks)
query <- GDCquery(project = "TCGA-LUAD",
                           data.category = "Gene expression",
                           data.type = "Gene expression quantification",
                           platform = "Illumina HiSeq",
                           experimental.strategy = "RNA-Seq",
              file.type  = "normalized_results",
                           legacy = TRUE)
GDCdownload(query, method = "api", files.per.chunk = 10)
data <- GDCprepare(query,summarizedExperiment=F)
```

2. Use *edgeR* [2] to further process the data to obtain counts per million (CPM) values per gene and sample and then use the marker genes *EPCAM*, *CLDN5*, *COL1A2*, and *PTPRC* to correlate sample-specific marker gene expression values to LMC proportions across the samples.

```
obj <- DGEList(data)
row.names(obj$samples) <- unlist(lapply(strsplit(row.names(obj$samples),"_"),
  function(x)x[3]))
colnames(obj$counts) <- unlist(lapply(strsplit(colnames(obj$counts),"_"),
  function(x)x[3]))
row.names(obj$samples) <- substr(row.names(obj$samples),1,16)
colnames(obj$counts) <- substr(colnames(obj$counts),1,16)
cpm.obj <- cpm(obj)
```

3. Plot each marker gene expression values per gene versus the LMC proportions.

```
load("FactorViz_outputs/medecom_set.RData")
props <- getProportions(medecom.set,K=7,lambda=0.001)
load("FactorViz_outputs/ann_S.RData")
colnames(props) <- substr(ann.S$Comment..TCGA.Barcode.,1,16)
marker.genes <- c("EPCAM","CLDN5","COL1A2","PTPRC")
in.exp <- colnames(cpm.obj) %in% colnames(props)
in.props <- colnames(props) %in% colnames(cpm.obj)
props <- props[,in.props]
cpm.obj <- cpm.obj[,in.exp]
cpm.obj <- cpm.obj[,colnames(props)]
row.names(cpm.obj) <- unlist(lapply(strsplit(row.names(cpm.obj),"[[:punct:]]"),
  function(x)x[1]))
cors.all <- sapply(marker.genes,function(marker){
  if(!marker %in% row.names(cpm.obj)){
    cors.gene <- NA
```

```
  }else{
    sel.exp <- cpm.obj[marker,]
    cors.gene <- apply(props,1,function(prop){
      cor(unlist(sel.exp),unlist(prop))
    })
  }
  cors.gene
})

cors.p.vals <- sapply(marker.genes,function(marker){
  if(!marker %in% row.names(cpm.obj)){
    cors.gene <- NA
  }else{
    sel.exp <- cpm.obj[marker,]
    cors.gene <- apply(props,1,function(prop){
      cor.test(unlist(sel.exp),unlist(prop))$p.value
    })
  }
  cors.gene
})
library(corrplot)
corrplot(cors.all,"ellipse")

plot.path <- "analysis/gene_expression/"
cors.all <- sapply(marker.genes,function(marker){
  if(!marker %in% row.names(cpm.obj)){
    cors.gene <- NA
  }else{
    sel.exp <- cpm.obj[marker,]
    for(j in 1:nrow(props)){
      prop <- props[j,]
      lmc <- paste0("LMC",j)
      to.plot <- data.frame(CPM=sel.exp,Proportion=prop)
      plot <- ggplot(to.plot,aes(x=Proportion,y=CPM))+geom_point(size=.1)+
        geom_smooth(method="lm",size=.5)+theme_bw()+
        theme(panel.grid=element_blank(),text=element_text(color="black",size=20),
          axis.ticks=element_line(size=0.5,color="black"),axis.ticks.length=unit(2,"mm"),
          axis.title=element_blank(),axis.text=element_blank())
      ggsave(file.path(plot.path,paste0(lmc,"_",marker,"_new.pdf")),
        plot,width=35,height=35,unit="mm")
    }
  }
})
```

#### Supplementary Tables

**Supplementary Table 1:** Overview of published DNA methylation based deconvolution tools. The methods are stratified according to the type and then ordered chronologically according to their date of publication.

| Tool | Type | Short description | Reference |
|---|---|---|---|
| *Houseman* | reference-based | The method employs constrained projection to infer proportions of reference profiles and was particularly developed for deconvolution of whole blood samples. | Houseman *et al.* [3], 2012 |
| *EpiDISH* | reference-based | *EpiDISH* is a reference-based method using robust partial correlations to compute proportions of reference profiles. The authors propose a method based on DNase hypersensitive sites to determine appropriate reference profiles. | Teschendorff *et al.* [4], 2017 |
| *hEpiDISH* | reference-base | *hEpiDISH* is an extension of *EpiDISH* that hierarchically performs deconvolution, and along with a new reference database, improves devonvolution results | Zheng *et al.* [5], 2018 |
| *Methyl-CIBERSORT* | reference-based | An extension of the *CIBERSORT* (Newman *et al.* [6], 2015) algorithm created for RNA-seq data that employs support vector regression (SVR) to estimate the proportions of given reference profiles across the samples. | Chakravarthy *et al.* [7], 2018 |
| *methylCC* | reference-based | *methylCC* uses latent components and a region-based, rather than an individual CpG-based, model to compute the proportions of given reference profiles independent of the technology (RRBS, WGBS, or BeadArray) used. | Hicks & Irizarry [8], 2019 |
| *IDOL* | selection of cell type markers | *IDOL* presents an improved strategy to determine cell-type specific marker CpGs, which improves deconvolution results | Salas *et al.* [9], 2018 |
| *FaST-LMM-EWASher* | confounding factor in EWAS | The *EWASher* approach is based on factored spectrally transformed linear mixed models to account for differences in cellular compositions in EWAS. | Zou *et al.* [10], 2014 |
| *ReFACTor* | confounding factor in EWAS | *ReFACTor* is based on Principal Component Analysis based on sites that are differentially methylated between cell types. The first few principal components are then used to adjust for cell type composition differences in EWAS. | Rahmani *et al.* [11], 2016 |
| *RefFreeCellMix* | reference-free | *RefFreeCellMix* from the *RefFreeEWAS* R-package uses non-negative matrix factorization (NMF) of the input DNA methylation matrix to compute a matrix of proportions and estimated reference profiles. | Houseman *et al.* [12], 2014 |
| *EDec* | reference-free | *EDec* is a two-step approach that combines reference-based and reference-free estimations using constrained matrix factorization. | Onuchic *et al.* [13], 2016 |
| *MeDeCom* | reference-free | *MeDeCom* uses regularized non-negative matrix factorization (NMF) of the input DNA methylation data matrix to create a matrix of proportions and of latent methylation components (LMCs). | Lutsik *et al.* [14], 2017 |
| *TCA* | reference-free | *TCA* uses tensor composition analysis to obtain sample-specific cell type profile estimates. In contrast to classical NMF, the method does not produce a single LMC matrix, but sample-specific LMCs using the same proportions matrix. | Rahmani *et al.* [15], 2019 |
| *CONFINED* | reference-free | *CONFINED* uses two matrices as input and employs canonical correlation analysis (CCA) to obtain purely biological sources of variations. | Thompson *et al.* [16], 2019 |
| *BayesCCE* | semi-reference-free | *BayesCCE* is a semi-supervised method to estimate proportions of different cell types that requires some prior knowledge on the cell-type composition of the studied tissue. | Rahmani *et al.* [17], 2018 |

**Supplementary Table 2:** Computational configurations in which software installation and the protocol have been tested. In case of an unexpected installation error, use the docker image available from https://hub.docker.com/r/mscherer/medecom.

| Type | Distribution | Version | R-version | Installation successful | Protocol tested | Comments |
|---|---|---|---|---|---|---|
| Linux | Debian | Wheezy (7) | R-3.5.2 | Yes | Yes | |
| | | | R-3.6.0 | Yes | Yes | |
| | | Jessie (8) | R-3.5.3 | Yes | Yes (reduced[1]) | |
| | | | R-3.6.1 | Yes | No | |
| | | | R-4.0 | Yes | No | |
| | | Buster (10) | R-3.5.2 | Yes | Yes (reduced) | |
| | Fedora | 28 | R-3.5.3 | Yes | No | |
| | | 31 | R-3.6.1 | No | Yes (reduced) | 'igraph' dependency fails to install |
| | CentOS | 8.0 | R-3.5.2 | Yes | Yes (reduced) | |
| | | | R-3.6.1 | Yes | Yes (reduced) | |
| | Ubuntu | 19 | R-3.6.1 | Yes | Yes (reduced) | |
| MacOS | | Mojave | R-3.5.1 | Yes | Yes (reduced) | binary release used |
| | | Catalina | R-3.6.0 | Yes | Yes (reduced) | |
| Windows | 10 | Pro | R-3.6.1 | No | Yes (reduced) | Use docker image |
| | 7 | Pro | R-3.6.1 | No | No | Docker is not available for Windows 7 |

[1] *In the reduced protocol, we executed preprocessing and a single MeDeCom run on a reduced dataset.*

**Supplementary Table 3:** Genomic annotations of the sites that had an absolute difference between LMC4 and the median of the other LMCs larger than 0.75. The distance corresponds to the distance of the CpG to the gene body of the closest gene (0 distance refers to sites located within the gene). CGI=CpG island, CTCF=CTCF binding site, ENSEMBL annotation=annotation according to the ENSEMBL regulatory build, proximal=proximal enhancer, TFBS=transcription factor binding site

| CpG ID | Chr | Start | End | Strand | CGI Relation | Difference | Closest gene | Closest gene (ENSEMBL) | Nearest gene distance | ENSEMBL annotation |
|---|---|---|---|---|---|---|---|---|---|---|
| cg00319661 | chr5 | 2632178 | 2632179 | + | Open Sea | -0.93 | C5orf38 | ENSG00000186493 | 120065 | |
| cg03415617 | chr16 | 34726856 | 34726857 | + | Open Sea | -0.922 | | ENSG00000260341 | 0 | CTCF |
| cg05789595 | chr6 | 56555274 | 56555275 | + | Open Sea | -0.864 | | ENSG00000231441 | 153524 | TSS |
| cg11006453 | chr8 | 141599185 | 141599186 | - | Open Sea | -0.851 | AGO2 | ENSG00000123908 | 0 | |
| cg08440178 | chr2 | 2737278 | 2737279 | + | Open Sea | -0.841 | MYT1L-AS1 | ENSG00000225619 | 406395 | |
| cg26992600 | chr14 | 37054509 | 37054510 | - | South Shore | 0.84 | NKX2-8 | ENSG00000136327 | 2696 | TFBS |
| cg25153741 | chr5 | 177913468 | 177913469 | + | Open Sea | -0.833 | RN7SL646P | ENSG00000242341 | 108248 | |
| cg13157980 | chr8 | 141599141 | 141599142 | - | Open Sea | -0.826 | AGO2 | ENSG00000123908 | 0 | |
| cg24066980 | chr2 | 1864151 | 1864152 | + | Open Sea | -0.821 | | ENSG00000232057 | 31711 | |
| cg23731089 | chr8 | 141599208 | 141599209 | - | Open Sea | -0.82 | AGO2 | ENSG00000123908 | 0 | |
| cg15616496 | chr17 | 73860607 | 73860608 | - | Open Sea | 0.816 | WBP2 | ENSG00000132471 | 8018 | |
| cg22986569 | chr5 | 2659008 | 2659009 | + | Open Sea | -0.811 | C5orf38 | ENSG00000186493 | 93235 | |
| cg26845946 | chr5 | 2137653 | 2137654 | - | South Shore | -0.803 | | ENSG00000248597 | 170384 | |
| cg03003434 | chr6 | 159141722 | 159141723 | - | Open Sea | -0.797 | AMZ2P2 | ENSG00000219249 | 5055 | |
| cg02896768 | chr1 | 154179996 | 154179997 | - | Open Sea | 0.793 | C1orf43 | ENSG00000143612 | 0 | |
| cg06255006 | chr16 | 86653215 | 86653216 | - | Open Sea | -0.786 | | ENSG00000260387 | 16717 | |
| cg11573608 | chr2 | 503193 | 503194 | - | Island | -0.779 | | ENSG00000223985 | 10537 | |
| cg16783478 | chr16 | 9943657 | 9943658 | - | Open Sea | -0.779 | GRIN2A | ENSG00000183454 | 0 | |
| cg06334134 | chr7 | 142986693 | 142986694 | + | South Shore | -0.779 | CASP2 | ENSG00000106144 | 0 | TSS |
| cg25453625 | chr7 | 4347751 | 4347752 | + | Island | -0.774 | SDK1 | ENSG00000146555 | 39118 | |
| cg03877767 | chr2 | 11680057 | 11680058 | + | Open Sea | 0.774 | GREB1 | ENSG00000196208 | 0 | TSS |
| cg14584961 | chr7 | 157533065 | 157533066 | + | Open Sea | -0.773 | | ENSG00000233038 | 114154 | TFBS |
| cg11761483 | chr17 | 70723386 | 70723387 | - | Open Sea | 0.772 | SLC39A11 | ENSG00000133195 | 0 | CTCF |
| cg02756683 | chr10 | 99449502 | 99449503 | - | South Shelf | -0.772 | AVPI1 | ENSG00000119986 | 2421 | TSS |
| cg17167920 | chr1 | 154127537 | 154127538 | + | Open Sea | 0.771 | UBAP2L | ENSG00000143569 | 65116 | |
| cg19075377 | chr13 | 112770169 | 112770170 | + | Open Sea | -0.77 | LINC00403 | ENSG00000224243 | 7839 | |
| cg26165146 | chr12 | 27484656 | 27484657 | + | North Shore | -0.765 | ARNTL2 | ENSG00000029153 | 1129 | TSS |
| cg05721751 | chr10 | 1707576 | 1707577 | + | Open Sea | -0.763 | ADARB2-AS1 | ENSG00000205696 | 108396 | |
| cg03945777 | chr7 | 157514049 | 157514050 | + | Open Sea | -0.759 | | ENSG00000222012 | 102591 | |
| cg20696049 | chr7 | 157551890 | 157551891 | + | South Shore | -0.759 | | ENSG00000233038 | 95329 | proximal |
| cg05726239 | chr6 | 107816677 | 107816678 | - | South Shelf | 0.757 | | ENSG00000234206 | 14328 | TSS |
| cg03262885 | chr7 | 157710179 | 157710180 | - | Open Sea | -0.757 | PTPRN2 | ENSG00000155093 | 0 | |
| cg14462553 | chr7 | 157444239 | 157444240 | - | South Shore | -0.757 | PTPRN2 | ENSG00000155093 | 0 | |
| cg06809074 | chr8 | 976415 | 976416 | - | Island | -0.756 | | ENSG00000254160 | 220372 | |
| cg26109981 | chr5 | 2175329 | 2175330 | + | Open Sea | -0.755 | | ENSG00000201026 | 9493 | |
| cg03540794 | chr5 | 2112109 | 2112110 | - | Island | -0.752 | | ENSG00000248597 | 144840 | |
| cg00327669 | chr5 | 1950782 | 1950783 | + | Island | -0.752 | | ENSG00000248994 | 0 | |
| cg26577252 | chr15 | 99212332 | 99212333 | + | Open Sea | -0.75 | IGF1R | ENSG00000140443 | 0 | |

### Supplementary Figures

**a** **Step I:** Start *FactorViz*

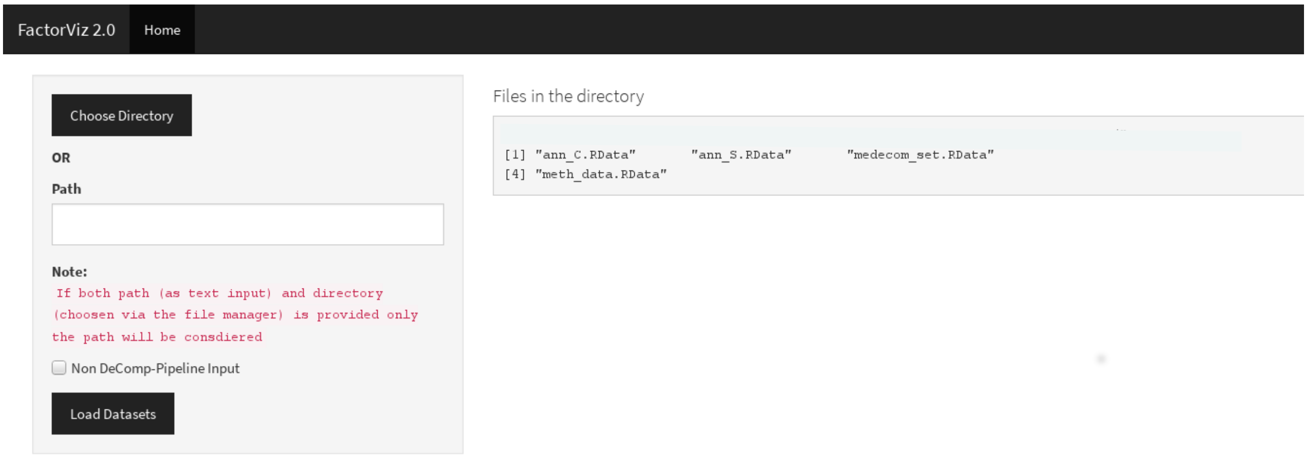


**b** **Step II:** Load *MeDeCom/DecompPipeline* output

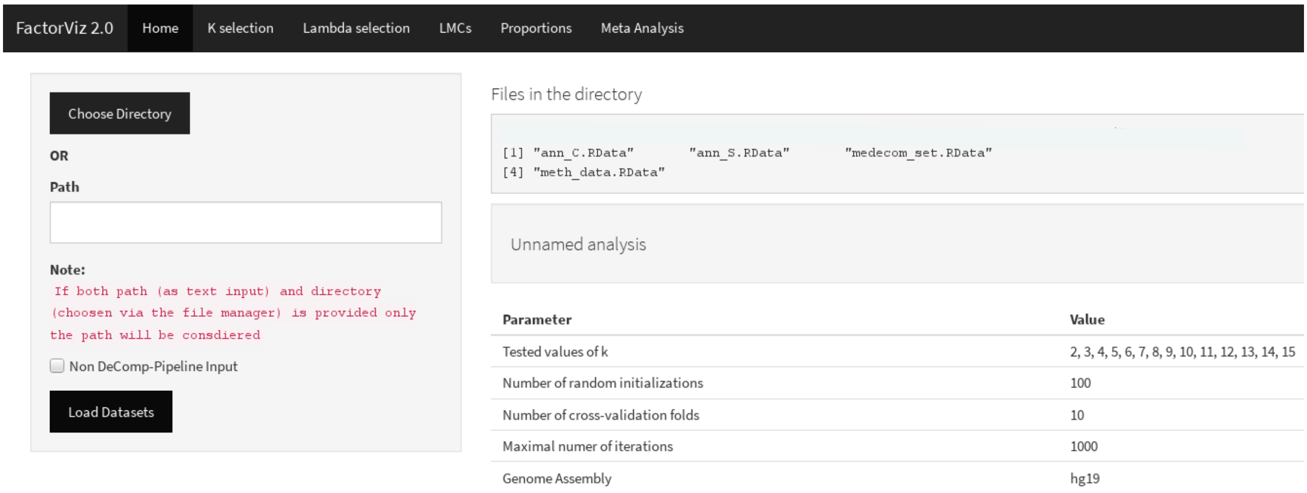

**c** **Step III:** Select number of components (K)

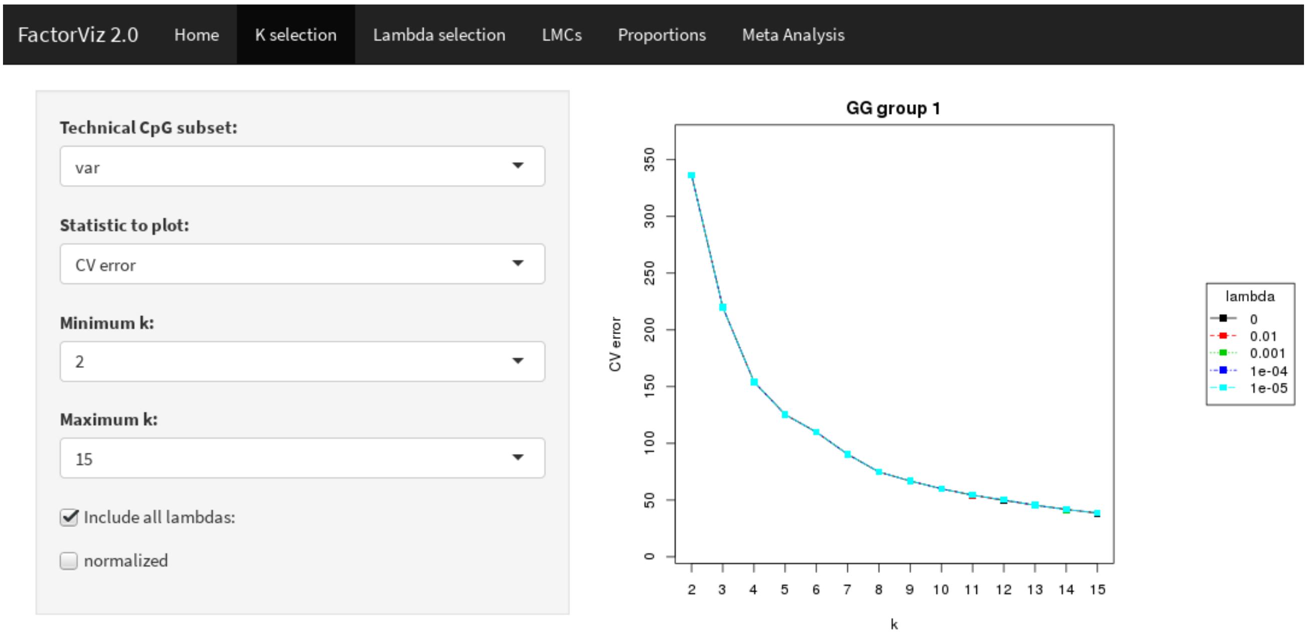


**d** **Step IV:** Select regularizer (λ)

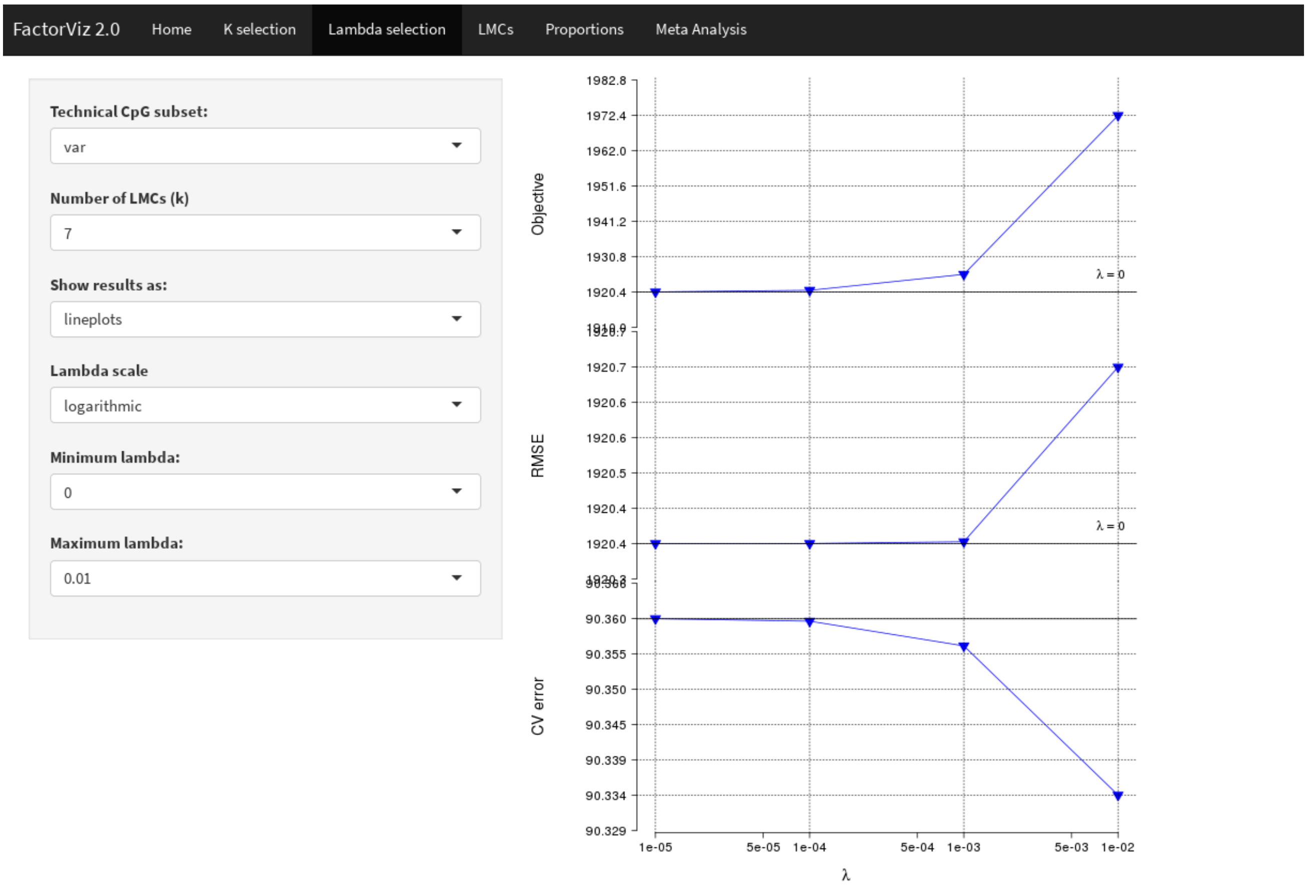

**e** **Step V:** Visualize proportions matrix (A)

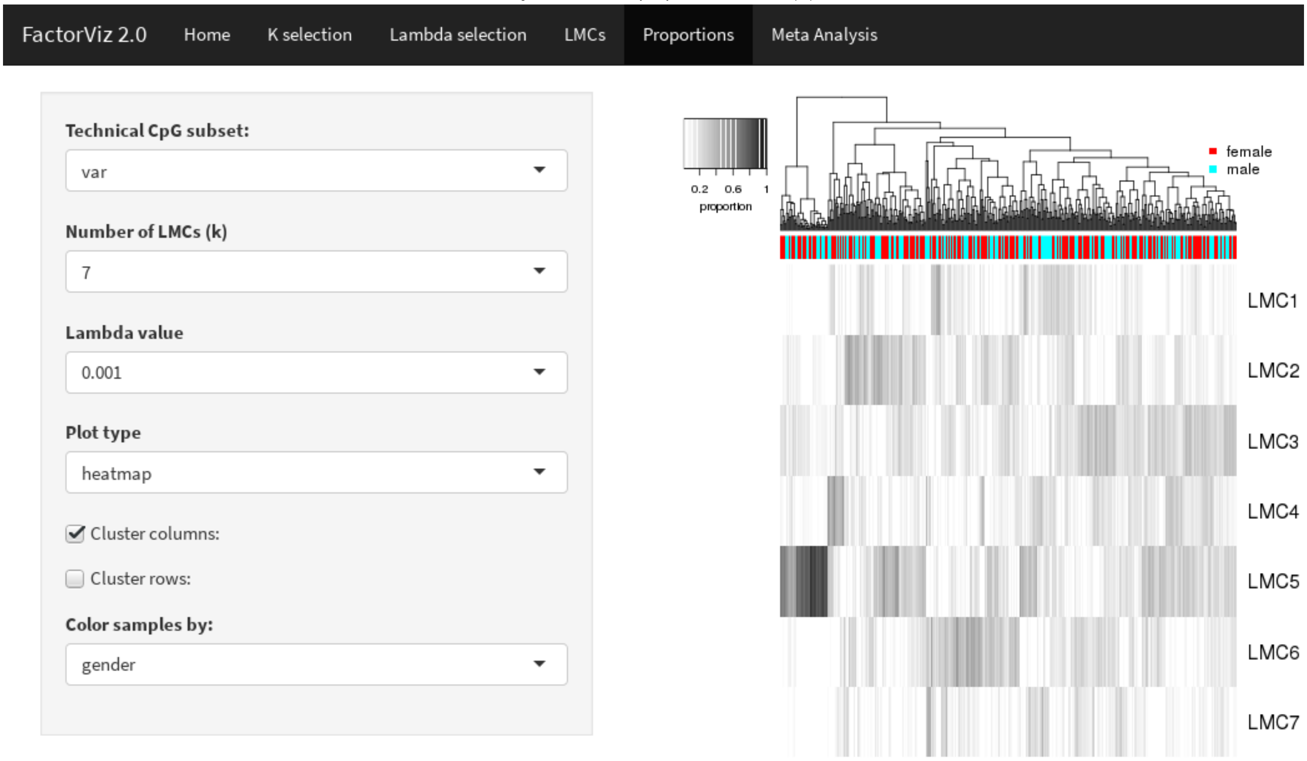


**f** **Step VI:** Associate proportions with phenotypes

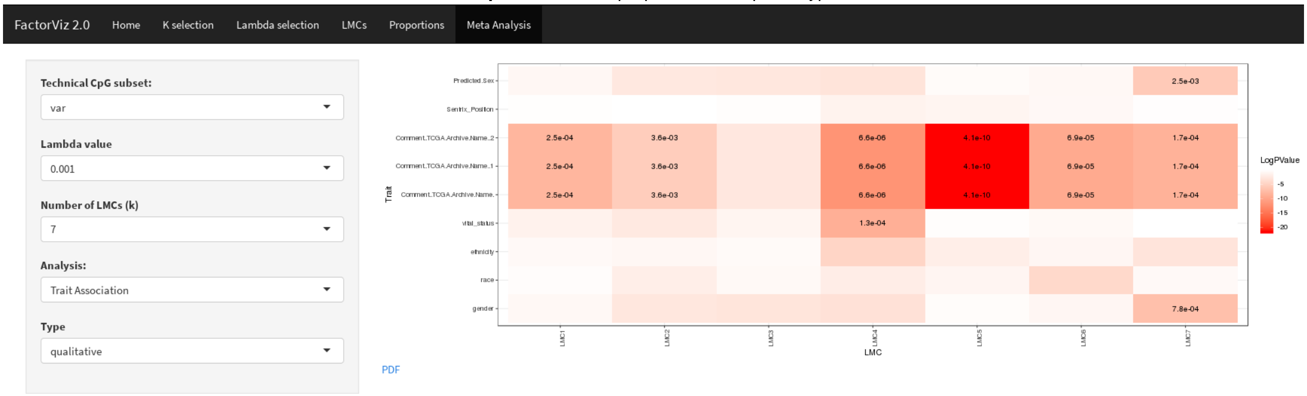

**g** **Step VII:** Visualize LMC matrix (T)

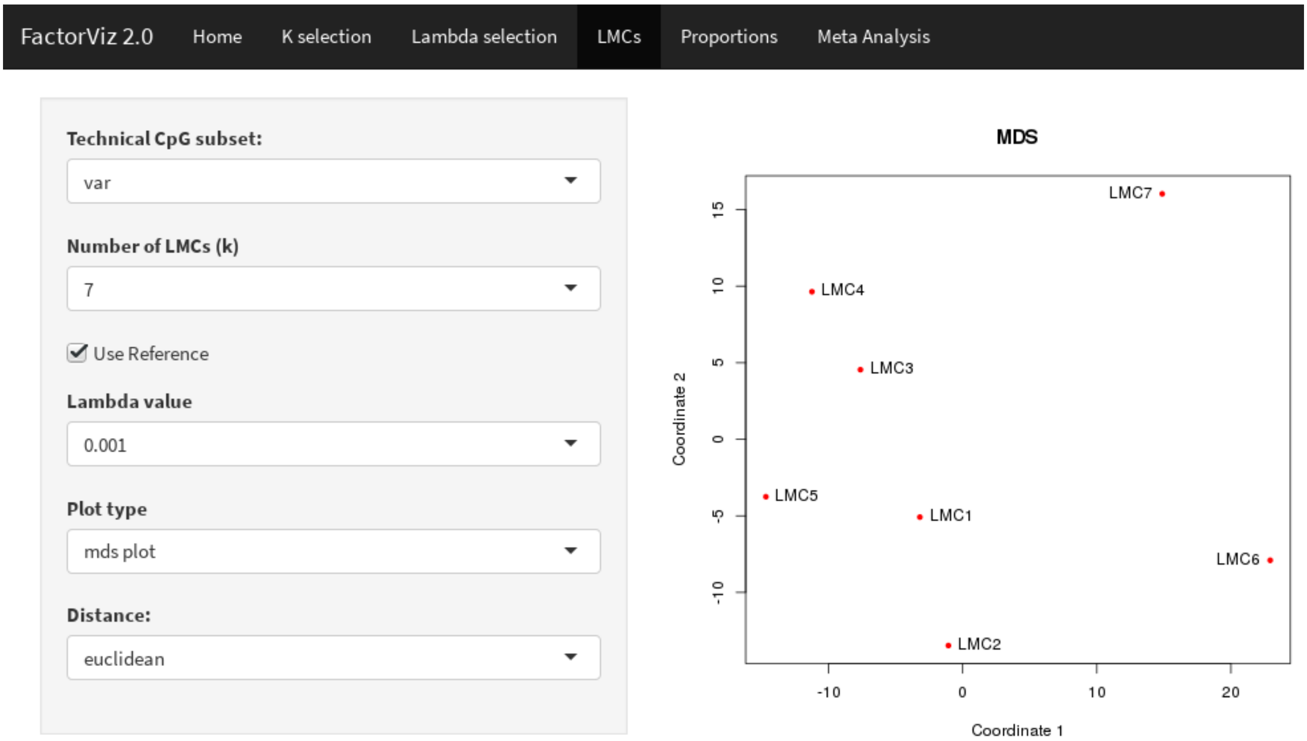


**h** **Step VIII:** Determine differential CpGs

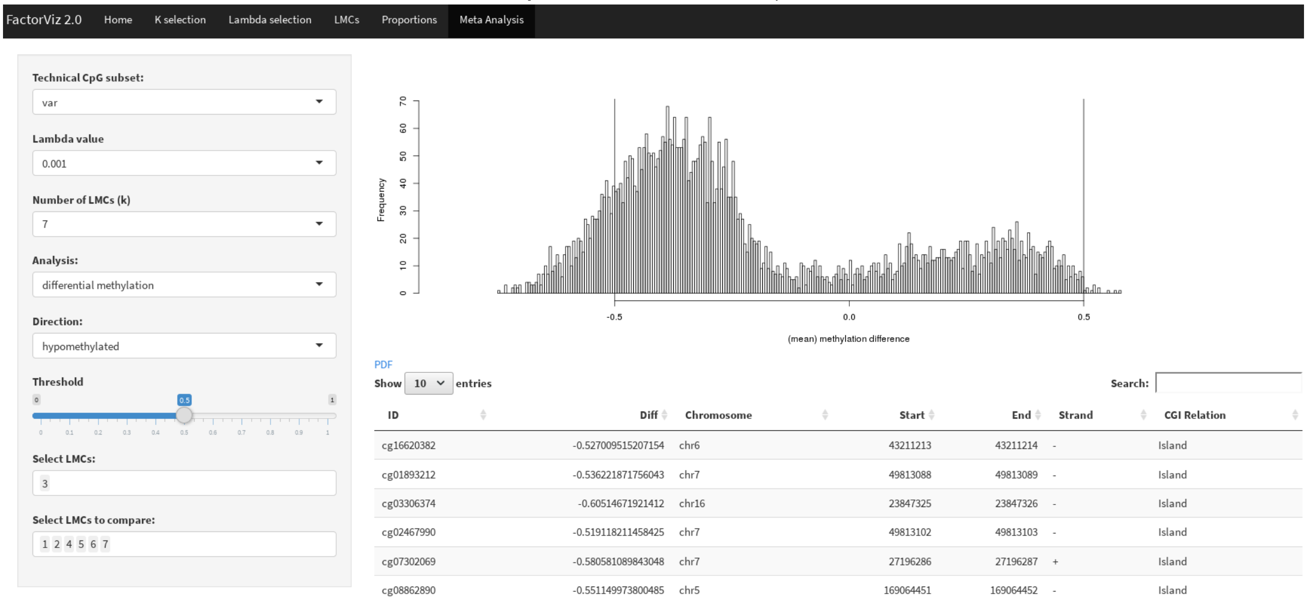

**i** **Step IX:** LMC GO enrichment analysis

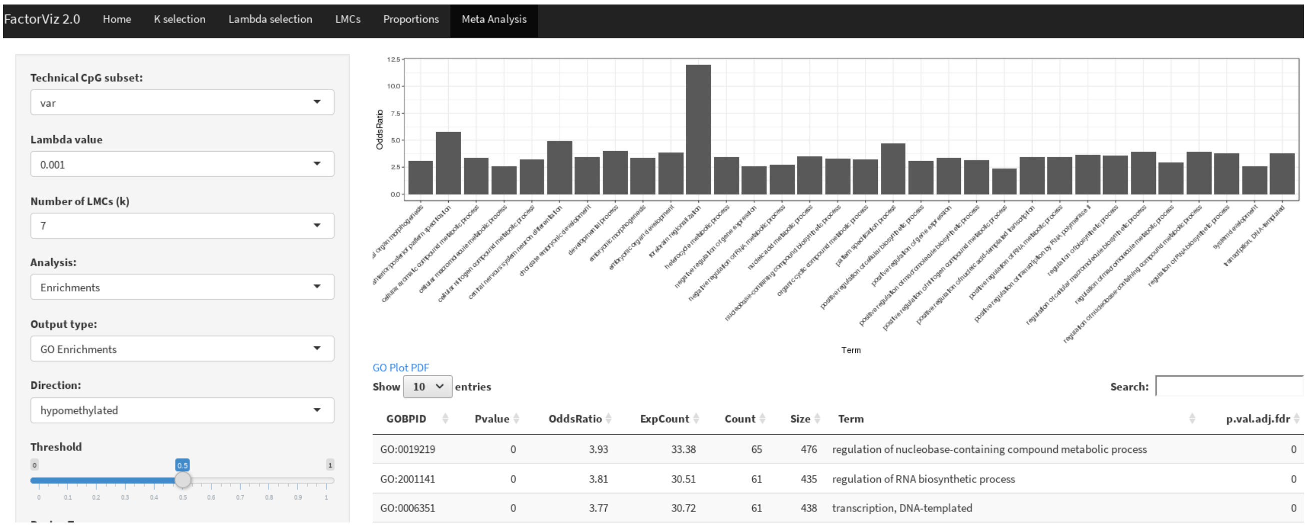


**j** **Step X:** LMC LOLA enrichment analysis

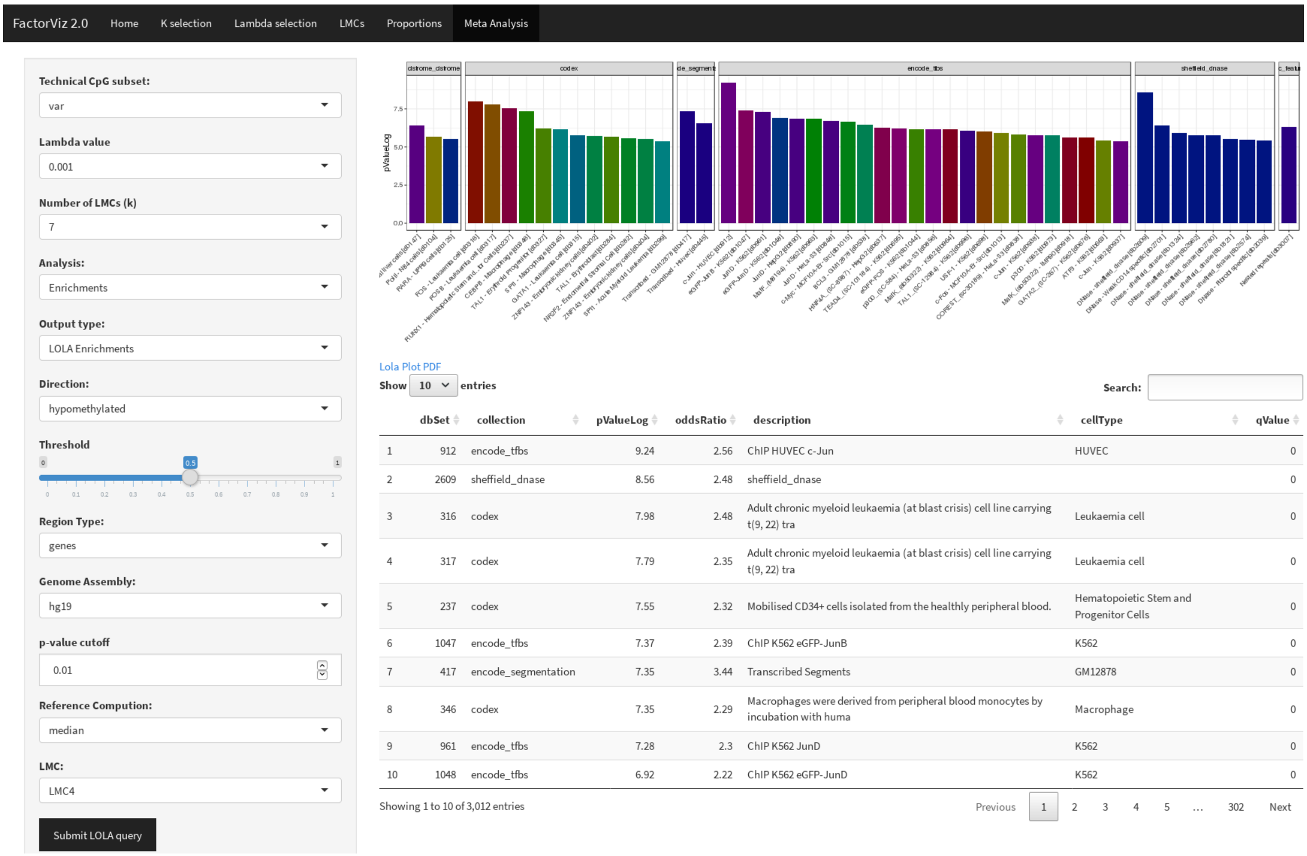


**Supplementary Fig. 1:** Interpreting *MeDeCom*'s results with *FactorViz*. For each of the steps, a screenshot of the *FactorViz* User Interface is shown for the TCGA LUAD dataset, and the ten performed steps are briefly described. **a, b** Specify the input, **c, d** Select the best parameters for the deconvolution, **e, f** Visualize proportion matrix and associate it with phenotypic traits, **g, h** Visualize LMCs matrix and determine differential CpGs, and **i, j** GO and LOLA enrichment analysis of differential CpGs.

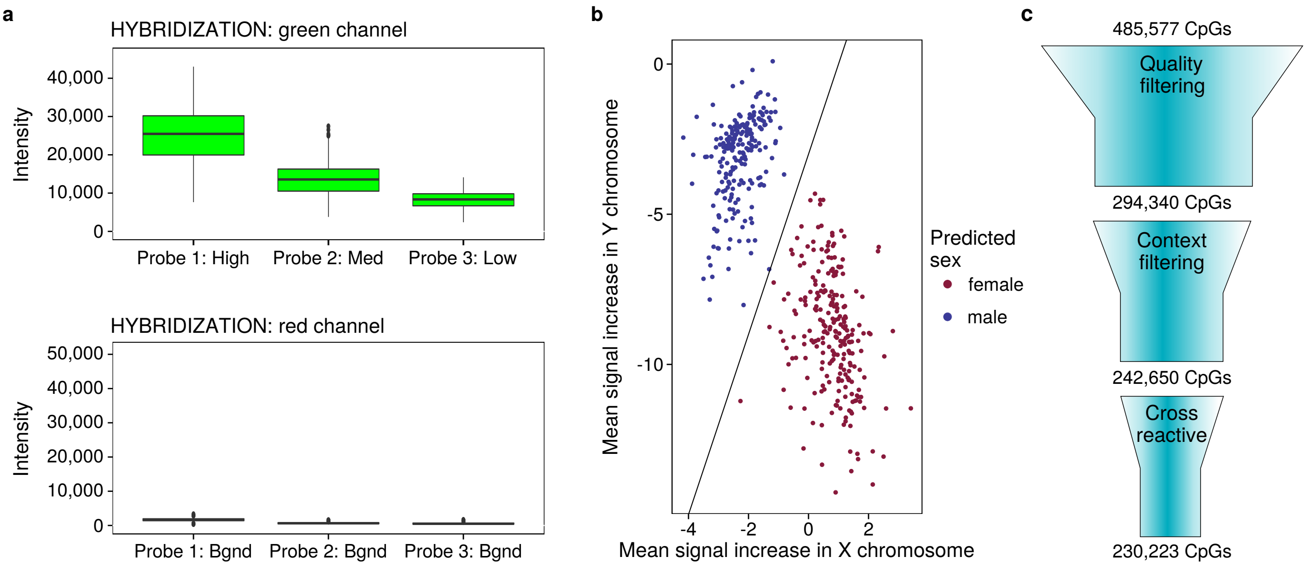


**Supplementary Fig. 2:** Quality control of TCGA data. **a** Boxplot for hybridization control probes for the green and the red channel, respectively. **b** Sex prediction based on the intensities of the probes on the sex chromosomes. A logistic regression classifier was employed to differentiate between female and male samples. **c** Outline of the CpG filtering procedure. The sites on the 450k array are filtered according to quality scores (coverage, overall intensity), genomic sequence context (SNPs, sex chromosomes), and cross-reactive sites are discarded.

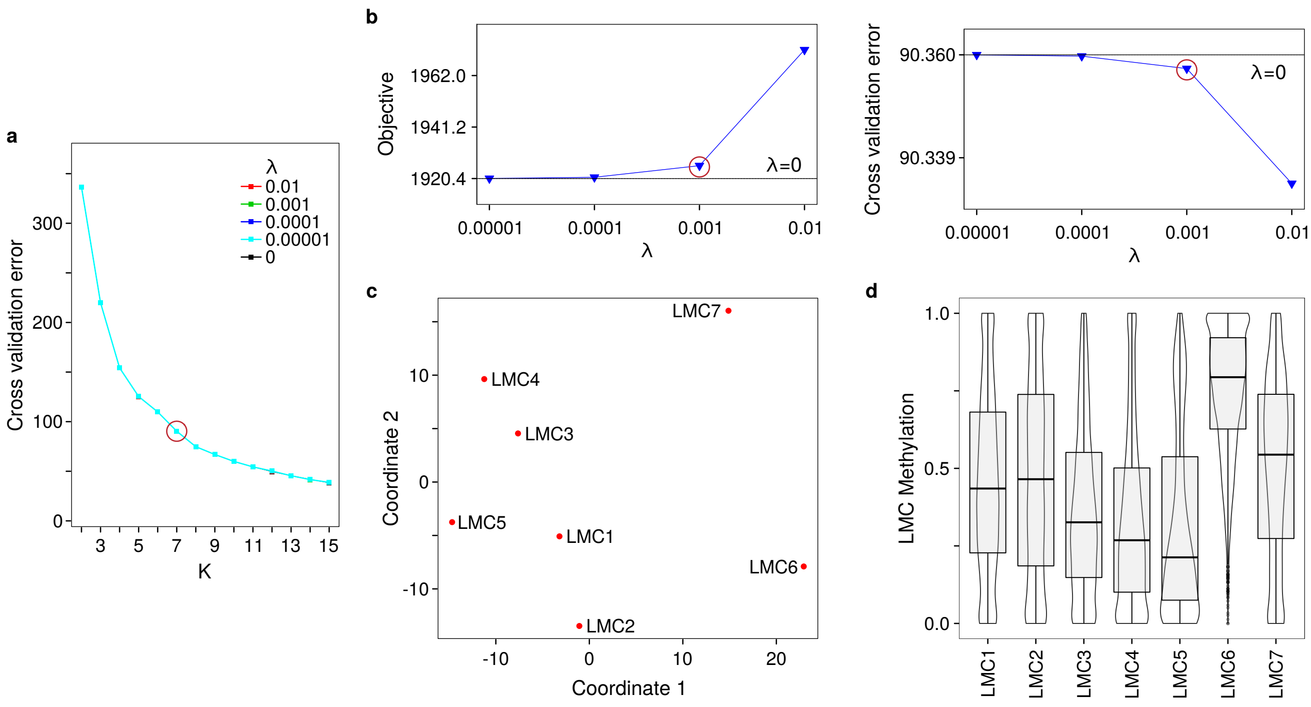


**Supplementary Fig. 3:** Selecting the number of components and the regularization parameter for *MeDeCom*. **a** Cross-validation error plotted against the number of latent components $K$ for different values of the regularization parameter $\lambda$. **b** Objective value and cross-validation error for different values of $\lambda$ after fixing the number of components to $7$. **c** Multidimensional scaling of the LMC data matrix after fixing the number of components to $7$ and the regularization parameter to $0.001$. Shown are the first two multidimensional components. **d** Violin plots of the LMC methylation matrix for the selected parameters.

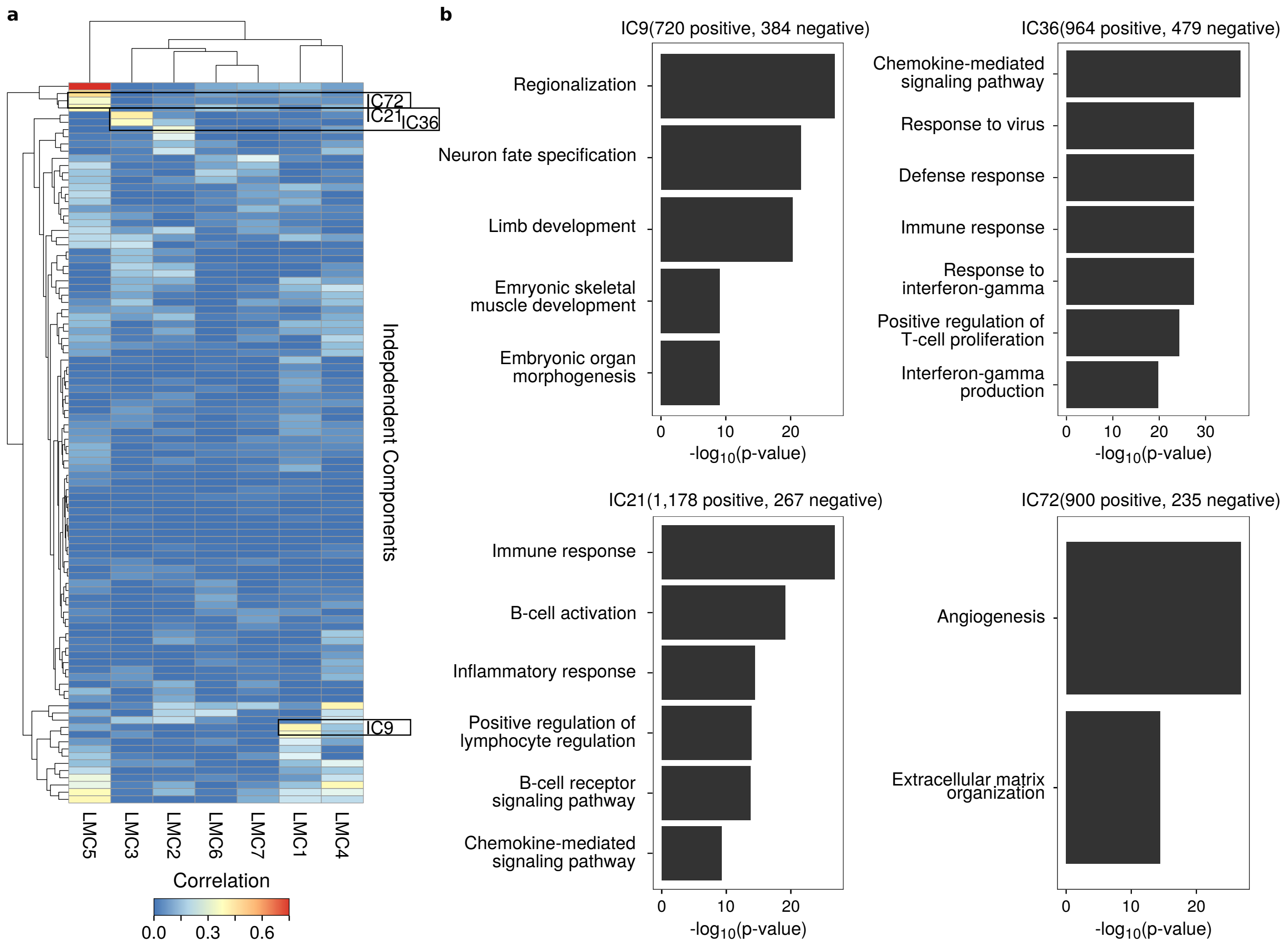


**Supplementary Fig. 4:** Comparing LMCs with independent components (ICs). **a** Correlation heatmap between the detected LMCs and the 100 detected independent components using ICA. Higher correlation is indicated by red and lower by blue colors. **b** GO enrichment analysis of the CpGs that contributed either positively or negatively (depicted in parentheses) to a particular independent component for IC9, IC21, IC36 and IC72.

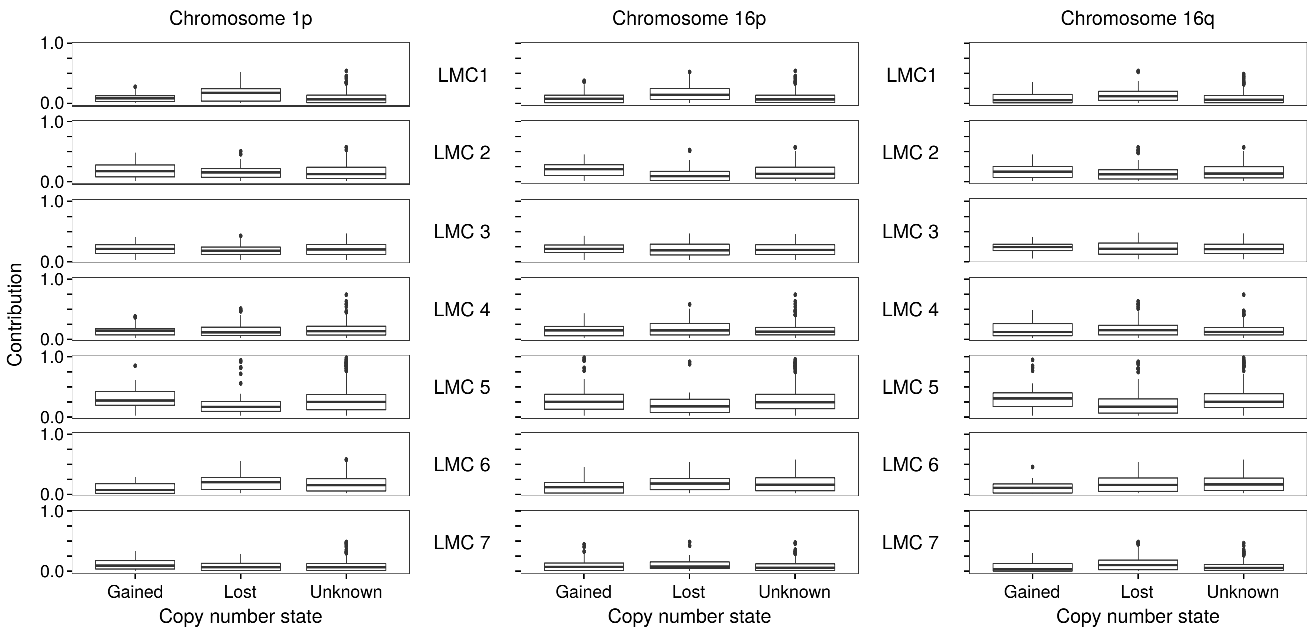


**Supplementary Fig. 5:** LMC (K=7, $\lambda$=0.001) contributions for different copy number states of different chromosomal parts in the TCGA LUAD dataset. The contributions have been stratified for each sample according to overall gain or loss of chromosomal parts. The copy number states were obtained from https://www.cbioportal.org/study/summary?id=luad_tcga_pan_can_atlas_2018 [18, 19].

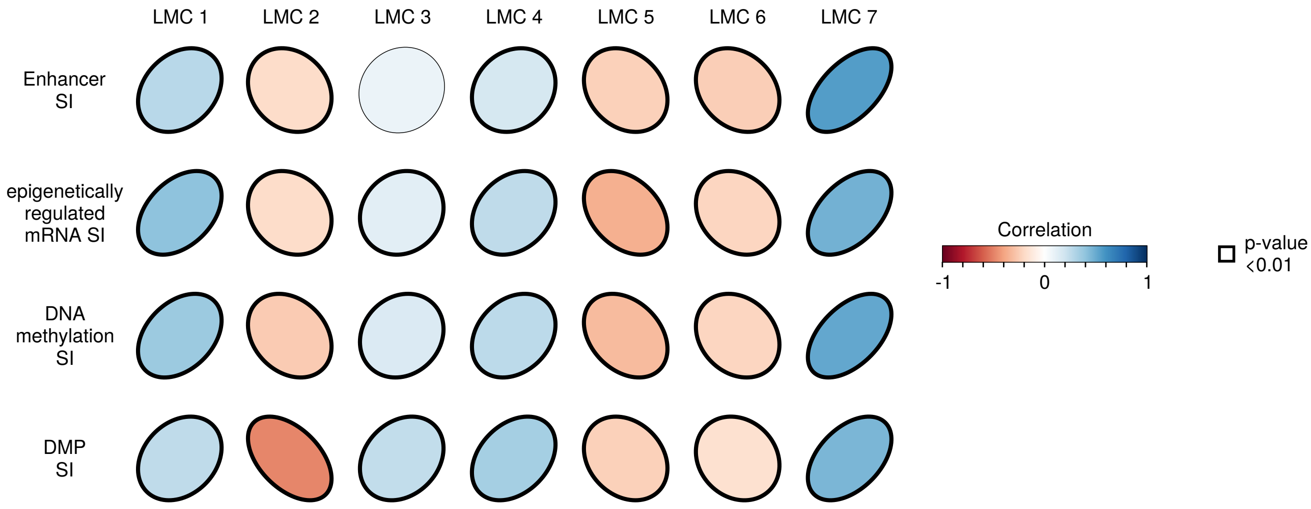


**Supplementary Fig. 6:** Pearson correlation between the different cancer stemness indices (SI) computed in Malta *et al.* [20] and the LMC proportions. The ellipses are directed towards the upper right for positive and to the lower right for negative correlations, respectively, while statistical significance is indicated by bold borders. DMP=differentially methylated probes

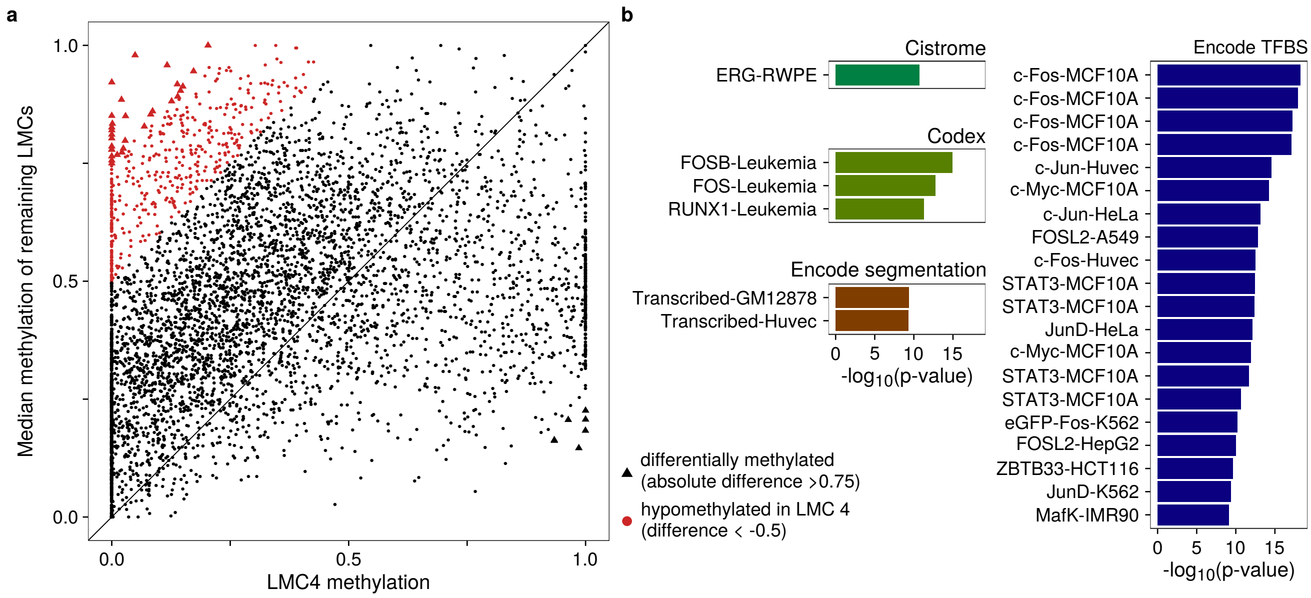


**Supplementary Fig. 7:** Differential analysis for LMC4. **a** Scatterplot between the methylation values of LMC4 (x-axis) and the median methylation values of the remaining six LMCs. Each point represents a CpG and points in red indicate the LMC-specific hypomethylated sites (difference less than 0.5), while the bold points represent those with an absolute difference larger than 0.75 (listed in **Supplementary Table 3**). **b** LOLA enrichment analysis of the LMC4-specific hypomethylated sites (the red points). Shown is the negative logarithm of the enrichment p-value.

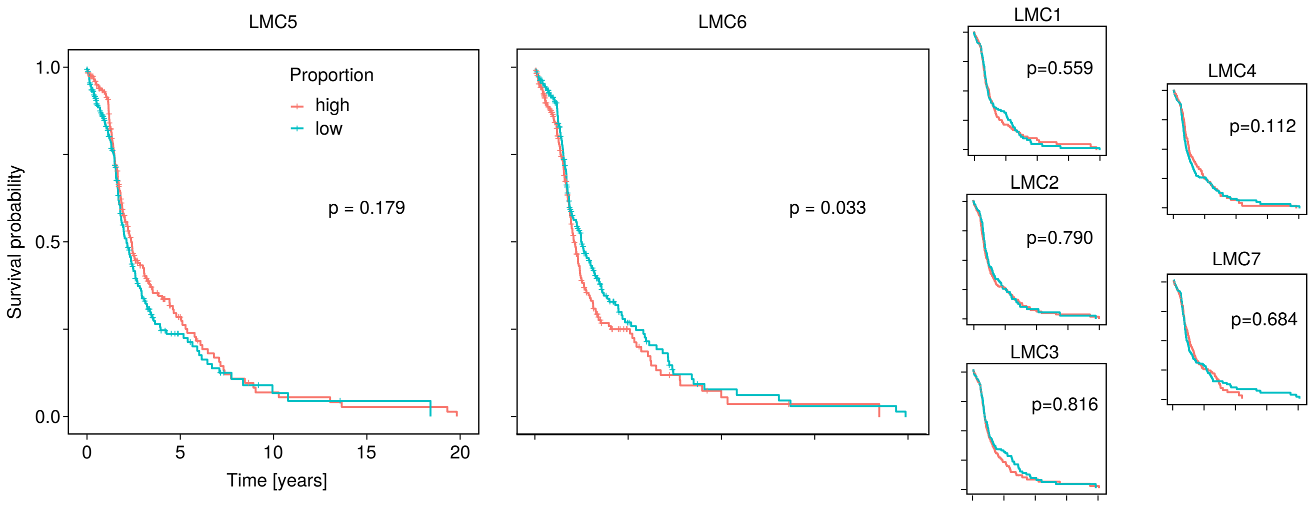


**Supplementary Fig. 8:** Survival analysis comparing different levels of LMC proportions. Shown are Kaplan-Meier curves, while samples were stratified according to the LMC proportions into two groups according to the median (high vs. low proportions). P-values were computed using the Cox proportional hazards model with the LMC proportions as input, and age, sex, and tumor stage as covariates [21].

### References


1. Colaprico, A. *et al.* TCGAbiolinks: An R/Bioconductor package for integrative analysis of TCGA data. *Nucleic Acids Res.* http://doi.org/10.1093/nar/gkv1507 (2015).
2. McCarthy *et al.* Differential expression analysis of multifactor RNA-Seq experiments with respect to biological variation. *Nucleic Acids Res.* **40,** 4288–4297 (2012).
3. Houseman, E. A. *et al.* DNA methylation arrays as surrogate measures of cell mixture distribution. *BMC Bioinf.* **13.** http://www.biomedcentral.com/1471-2105/13/86 (2012).
4. Teschendorff, A. E., Breeze, C. E., Zheng, S. C. & Beck, S. A comparison of reference-based algorithms for correcting cell-type heterogeneity in Epigenome-Wide Association Studies. *BMC Bioinf.* **18,** 105. ISSN: 1471-2105. http://www.ncbi.nlm.nih.gov/pubmed/28193155http://www.pubmedcentral.nih.gov/articlerender.fcgi?artid=PMC5307731 (2017).
5. Zheng, S. C. *et al.* A novel cell-type deconvolution algorithm reveals substantial contamination by immune cells in saliva , buccal and cervix. *Epigenomics* **10,** 925–940. ISSN: 1750-192X (2018).
6. Newman, A. M. *et al.* Robust enumeration of cell subsets from tissue expression profiles. *Nat. Methods* **12,** 453–457. http://www.ncbi.nlm.nih.gov/pubmed/25822800 (2015).
7. Chakravarthy, A. *et al.* Pan-cancer deconvolution of tumour composition using DNA methylation. *Nat. Commun.* **9** (2018).
8. Hicks, S. C. & Irizarry, R. A. methylCC: technology-independent estimation of cell type composition using differentially methylated regions. *Genome Biol.* **20,** 261. ISSN: 1474-760X. https://www.biorxiv.org/content/early/2017/11/03/213769https://genomebiology.biomedcentral.com/articles/10.1186/s13059-019-1827-8 (2019).
9. Salas, L. A. *et al.* An optimized library for reference-based deconvolution of whole-blood biospecimens assayed using the Illumina HumanMethylationEPIC BeadArray. *Genome Biol.* **19,** 64. ISSN: 1474-760X. https://www.ncbi.nlm.nih.gov/geo/query/acc.cgi?acc=GSE110554https://genomebiology.biomedcentral.com/articles/10.1186/s13059-018-1448-7 (2018).
10. Zou, J., Lippert, C., Heckerman, D., Aryee, M. & Listgarten, J. Epigenome-wide association studies without the need for cell-type composition. *Nat. Methods* **11,** 309–311. ISSN: 15487105 (2014).
11. Rahmani, E. *et al.* Sparse PCA corrects for cell type heterogeneity in epigenome-wide association studies. *Nat. Methods* **13,** 443–445. http://www.nature.com/doifinder/10.1038/nmeth.3809 (2016).
12. Houseman, E. A., Molitor, J. & Marsit, C. J. Reference-free cell mixture adjustments in analysis of DNA methylation data. *Bioinformatics* **30,** 1431–1439. ISSN: 1367-4803. https://academic.oup.com/bioinformatics/article-lookup/doi/10.1093/bioinformatics/btu029 (2014).
13. Onuchic, V. *et al.* Epigenomic Deconvolution of Breast Tumors Reveals Metabolic Coupling between Constituent Cell Types. *Cell Reports* **17,** 2075–2086. ISSN: 22111247. arXiv: 15334406. http://dx.doi.org/10.1016/j.celrep.2016.10.057 (2016).

14. Lutsik, P. *et al.* MeDeCom: discovery and quantification of latent components of heterogeneous methylomes. *Genome Biol.* **18,** 55. http://genomebiology.biomedcentral.com/articles/10.1186/s13059-017-1182-6 (2017).
15. Rahmani, E. *et al.* Cell-type-specific resolution epigenetics without the need for cell sorting or single-cell biology. *Nat. Commun.* **10** (2019).
16. Thompson, M., Chen, Z. J., Rahmani, E. & Halperin, E. CONFINED: Distinguishing biological from technical sources of variation by leveraging multiple methylation datasets. *Genome Biol.* **20,** 1–15. ISSN: 1474760X (2019).
17. Rahmani, E. *et al.* BayesCCE: a Bayesian framework for estimating cell-type composition from DNA methylation without the need for methylation reference. *Genome Biol.* **19,** 1–18. ISSN: 1474760X (2018).
18. Cerami, E. *et al.* The cBio Cancer Genomics Portal: An open platform for exploring multidimensional cancer genomics data. *Cancer Discovery* **2,** 401–404. ISSN: 21598274 (2012).
19. Gao, J. *et al.* Integrative Analysis of Complex Cancer Genomics and Clinical Profiles Using the cBioPortal. *Sci. Signal.* **6,** pl1–pl1. ISSN: 1945-0877. http://stke.sciencemag.org/cgi/doi/10.1126/scisignal.2004088 (2013).
20. Malta, T. M. *et al.* Machine Learning Identifies Stemness Features Associated with Oncogenic Dedifferentiation. *Cell* **173,** 338–354.e15. ISSN: 10974172 (2018).
21. Therneau, T. M. *A Package for Survival Analysis in S* version 2.38 (2015). https://CRAN.R-project.org/package=survival.
